# Mapping mesenchymal diversity in the developing human intestine and organoids

**DOI:** 10.1101/2025.07.22.665939

**Authors:** Kelli F. Johnson, Xiangning Dong, Yu-Hwai Tsai, Angeline Wu, Sydney G. Clark, Sha Huang, Rachel K. Zwick, Ian Glass, Katherine D. Walton, Ophir D. Klein, Jason R. Spence

**Affiliations:** Department of Internal Medicine, Division of Gastroenterology, University of Michigan Medical School, Ann Arbor, MI 48109, USA; Department of Cellular and Molecular Biology, University of Michigan Medical School, Ann Arbor, MI 48109, USA; Department of Cell Biology, NYU Grossman School of Medicine, New York, NY, USA; Regenerative Medicine Institute, NYU Grossman School of Medicine, New York, NY, USA; Department of Pediatrics, Genetic Medicine, University of Washington, Seattle, WA 98195, USA; Program in Craniofacial Biology and Department of Orofacial Sciences, University of California, San Francisco, San Francisco, CA, USA; Department of Pediatrics, Cedars-Sinai Guerin Children’s, Los Angeles, CA, USA; Department of Cell and Developmental Biology, University of Michigan Medical School, Ann Arbor, MI 48109, USA; Department of Biomedical Engineering, University of Michigan and University of Michigan College of Engineering, Ann Arbor, MI, USA

**Keywords:** development, mesenchyme, fibroblast, single-cell sequencing, spatial transcriptomics, Xenium, human, intestine

## Abstract

The organization of diverse mesenchymal populations during human intestinal development is critical for tissue architecture and function yet remains poorly defined. To construct a comprehensive, tissue-scale map of the developing human small intestine, we leveraged single-cell RNA-sequencing data to build a custom Xenium spatial transcriptomics gene panel covering the diversity of cell types in the human intestine. Analysis was focused on the developing mesenchyme populations (also referred to as fibroblasts or stroma) given the lack of spatiotemporal information about these cell populations. We defined 5 broad mesenchymal populations occupying discrete anatomical locations within the lamina propria and submucosa – the subepithelial cells (SEC), lamina propria fibroblasts (LPF), submucosal fibroblasts (SMF), smooth muscle cells (SMC), and *CXCL13*+ fibroblasts. Our data reveal dynamic spatial remodeling of fibroblast communities during development and establish molecular markers to distinguish these populations. We leverage this high-resolution atlas to benchmark pluripotent stem cell-derived human intestinal organoids and to demonstrate how this foundational resource can be used to dissect intestinal stromal signaling in a spatial manner, with broad implications for modeling development, regeneration, and disease.

**Graphical Abstract:** 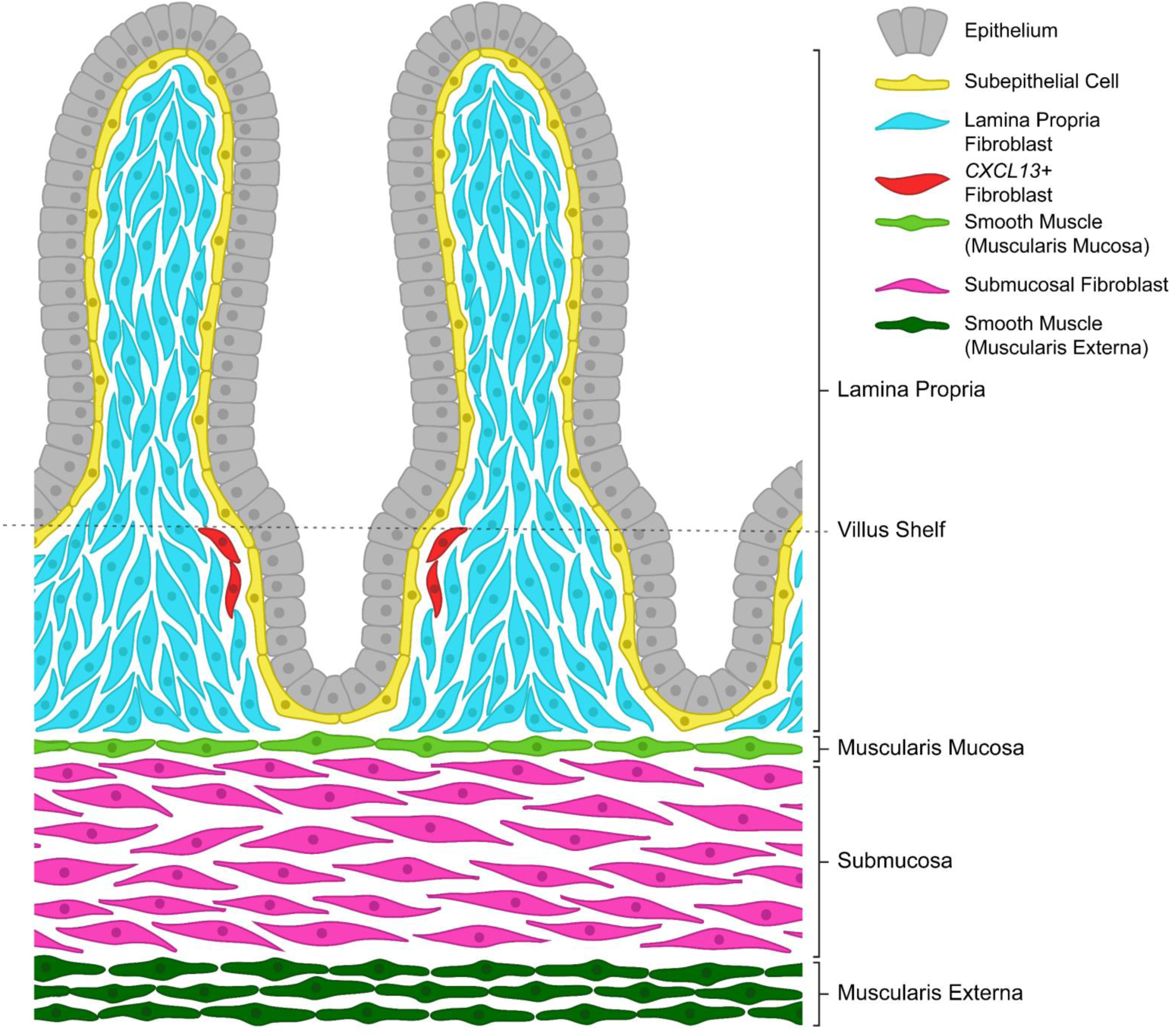

*Highlights:*

- A spatial atlas of fibroblast heterogeneity in the human intestine
- Populations include subepithelial, submucosal, lamina propria, and *CXCL13*+ fibroblasts
- Fibroblast position in the developing intestine is maintained into adulthood
- Organoids largely recreate the spatial organization of the human intestine

## Introduction

From the characteristic crypt-villus axis that enables a large surface area for nutrient absorption to the distinct muscle layers that coordinate with the enteric nervous system to generate peristaltic and propulsive motion, the precise organization of the small intestine is essential for successful function. The cellular heterogeneity that contributes to this architecture both in the adult and developing human intestine have been the focus of many recent studies utilizing single cell approaches to understand development, homeostasis, and disease^1–15^. While tissue-scale processes and features such as villus morphogenesis and the cellular composition of the intestine are well established for several lineages such as the intestinal epithelium, musculature, and enteric nervous system^1,6,9,14,16–26^, the mesoderm-derived fibroblast populations (also referred to as mesenchyme and/or stroma) remain poorly defined.

In mice, recent efforts have led to a revisiting of historical terms such as ‘fibroblast’, ‘myofibroblast’, and ‘intestinal subepithelial myofibroblast’ (ISEMF)^27–38^ based on molecular identity, cellular function, and cellular localization^39,40^. For example, recent studies have described the subepithelial ‘telocytes’ or ‘subepithelial myofibroblasts’ (SEMFs) as cells defined by high levels of *Pdgfra* and *Foxl1*, as well as two populations of *Pdgfra*-low expressing cells, the *Cd81*+/*Grem1*+ ‘trophocytes’ and a *Cd34*+/*Pdpn*+ “*Pdgfra*-low stroma”^29,41–43^. Another recent study demonstrated that distinct regional mesenchymal populations line the developing mouse foregut and contribute to patterning the endoderm of the GI tract^44^, indicating functional distinctions for transcriptionally unique populations based on early localization. However, whether spatially distinct fibroblast populations also support development and patterning of the midgut is unknown. Further, much of this work toward characterizing intestinal fibroblast domains was conducted in mice. The question of how much is relevant to human development remains unaddressed.

These studies have begun to advance our understanding of the cellular heterogeneity within the mouse intestinal subepithelial compartment, but an overarching atlas of the distinct identities and positions of each fibroblast subtype relative to all other cell types has not been defined in either the mouse or the human^29^. Here, we aimed to systematically define the molecular and spatiotemporal heterogeneity of the developing human intestine, with an emphasis on the poorly defined fibroblasts. Our results showed that the intestinal fibroblast population is made up of transcriptionally distinct sub-populations that occupy unique spatial domains within the developing human small intestine.

In order to address the spatial localization and cellular heterogeneity, we integrated publicly available single cell RNA-sequencing (scRNA-seq) datasets from multiple studies spanning 5-21 weeks post-conception^1,5,45,46^. From these data, we defined sets of highly enriched genes that allowed us to identify cell types of interest within the dataset using a molecular signature and then used this information to generate a 10X Xenium custom probe set^47–51^. Within the probe set, we also included published and well-established intestinal marker genes, as well as genes related to well-known intestinal signaling and structural pathways. Using this probe set, we interrogated a total of seven different developing human intestines spanning 7-22 weeks post-conception, including all main regions of the intestinal tract (duodenum, jejunum, ileum, colon).

Xenium imaging and cell segmentation produced a cellular-resolution atlas of the intestinal architecture. We visualized all major cell classes (epithelium, fibroblasts, smooth muscle, endothelium, enteric nervous system, and immune) and their organization. To further interrogate the intestinal fibroblasts, we examined all non-muscle fibroblasts in both the scRNA-seq and Xenium data. In addition to smooth muscle cell (SMC) populations, we identified 4 broad mesenchymal fibroblast sub-populations that occupy discrete anatomical locations within the lamina propria and submucosa, including subepithelial cells (SECs)^1,7^, lamina propria fibroblasts (LPFs), submucosal fibroblasts (SMFs), and *CXCL13*+ fibroblasts^7^. Our data reveal spatial localization of fibroblast populations during development and establish molecular markers to distinguish these populations.

Finally, we used spatial transcriptomics to interrogate two different iPSC-derived human intestinal organoid models following transplantation into mice (transplanted HIOs; tHIOs)^52–55^. Using the human intestinal spatial atlas as a framework for understanding tHIOs, we identified subtle changes within HIOs derived by different methodologies, suggesting that not only is accurately representing cellular heterogeneity an important measure of HIO fidelity to the *in vivo* human intestine, but so is the spatial organization of cellular populations within the HIOs.

Altogether, our work provides a tissue-scale, spatially resolved blueprint of the developing human intestine with a focus on the fibroblast/mesenchyme populations, establishing molecular signatures that allow us to identify fibroblast identities and track their changes across development. By using this atlas to assess the spatial organization of tHIOs, we reveal that fidelity must be judged not only by cell-type composition but also by the precise topography of cells, and we anticipate that this will sharpen organoid design, disease modeling, and therapeutic screening. Beyond filling a critical knowledge gap, the resource we provide transforms our current understanding of the transcriptomic landscape of the gut into a three-dimensional coordinate system that developmental biologists, tissue engineers, and gastroenterologists can apply with high precision.

## Results

### Spatial transcriptomics identifies major cell classes in the developing human intestine

Based on our previously published scRNA-seq data^1^ (**Figure 1A-B, Supplemental Table 1**), established marker genes, and genes of interest, we designed a custom Xenium panel of 292 genes to identify the cell types/states found in the developing human small intestine^1,3,30,31^ (**Figure 1C-D, Supplemental Table 2,** and **Methods**).

**Figure 1:**
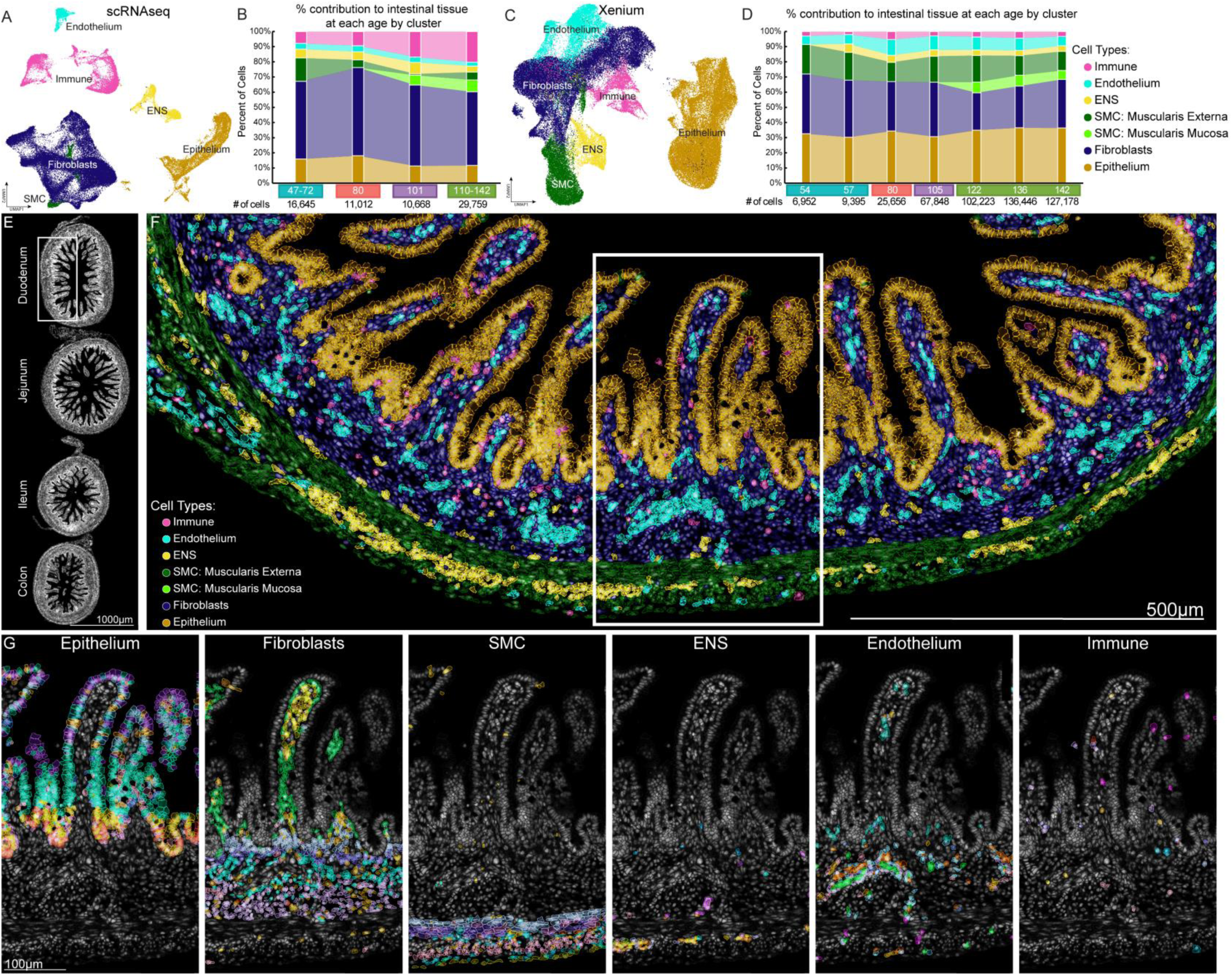
Comparison of major cell type class composition between scRNAseq and Xenium datasets describing the human fetal small intestine. UMAP projections summarizing the major cell classes observed in human fetal small intestine as determined by **A)** scRNAseq and **C)** Xenium spatial transcriptomics. Stacked bar graphs describing the cell population composition of the human fetal small intestine at various time points through development (47-142 DPC) for either the **B)** scRNAseq dataset or **D)** Xenium dataset. Sample labels below the graphs in **B and D** are colored to indicate the developmental stage (DPC) each sample falls within. The number of cells included in each sample is indicated below the sample age. **E-G** depict the 105 DPC human fetal duodenum as a representative Xenium image set example. **E)** The DAPI image of the complete Xenium 105 DPC sample. Scalebar = 1000µm. The white box outlines the region depicted in **F**. **F)** A digital cell mask depicting the distribution of the major cell classes annotated in **C**. Scalebar = 500µm. The white box indicates the region depicted in **G**. **G)** An example of cell sub-type distribution within each major cell class identified using the Xenium panel visualized with each color representing distinct sub-types: **G1)** the epithelium sub-types, **G2)** the fibroblast sub-types, **G3)** the SMC-related sub-types, **G4)** the ENS sub-types, **G5)** the endothelium sub-types, and **G6**) the immune sub-types. Scalebar = 100µm. Cell populations are color matched in **A-F**; epithelium (gold), fibroblasts (navy), muscularis mucosa (MM, light green), other SMCs (dark green), ENS (yellow), endothelium (cyan), and immune (pink). DAPI staining of cell nuclei depicted in grey.

Among the major cell classes, we used well known markers, for example, *CDH1* for the epithelium, *VIM* for the mesenchyme, *PECAM1* for the endothelium, *PTPRC (CD45)* for the immune cells, and *TUBB3* for the enteric nervous system (ENS) (**Supplemental Table 3**). We also included markers that would identify sub-classes of cells, with a focus on interrogating the different types of mesenchymal fibroblast populations identified in scRNA-seq data^1^. Finally, we carried out Xenium spatial transcriptomics on formalin fixed, paraffin embedded (FFPE) archival tissue samples across the intestine (**Figure 1E** - duodenum, jejunum, ileum, colon) and ranging from 54 days post-conception (DPC) to 142 DPC, roughly matching the stages included in the scRNA-seq dataset (**Supplemental Table 1**). In total, we visualized n=7 different biological specimens.

While we imaged sections from all regions of the intestine, our analysis focused on the small intestine which included a total of 475,698 cells across all stages (**Figure 1C-D**; **Methods**). The distal ileum (when included in the sample imaging) and the colon were excluded from further analysis due to significant transcriptomic differences between the small intestine and colon^40^.

We carried out spatial transcriptomics on two batches of samples. Due to improvements to the Xenium technology, samples run in the second batch were imaged with the Xenium *In Situ* Multimodal Cell Segmentation kit (n=5 samples), which was not available during our first batch of sequencing (n=3 samples; 57, 80, and 142 DPC). The kit included a mix of probes and antibodies for cell borders and cell interiors to more accurately define the cell bodies of as many cell types as possible during digital segmentation (**Supplemental Figure 1C**). Digital segmentation of the Xenium data was performed using the Xenium Ranger analysis pipeline and visualized with Xenium Explorer (10X Genomics, v2.0.0 and later) (**Supplemental Figure 1D-E**). Xenium data acquired without the use of the segmentation kit (Batch 1: samples 57, 80, and 142 DPC) were segmented using the internal segmentation algorithm of Xenium Explorer.

The resulting segmentation maps from either method were used in conjunction with the (X, Y) coordinates of every positive probe location for clustering and analysis of the Xenium data, including graph-based clustering (**Supplemental Figure 1E**).

### Xenium captures the diversity of cell types identified in scRNA-seq data

Following cell segmentation, the cell identification (cell-id) and associated transcript data for the small intestinal regions (duodenum, jejunum, ileum) were extracted and imported into Seurat^56^ (**Figure 1C-D**). Using an analysis pipeline (**Methods**) similar to the one applied to scRNA-seq data, we generated UMAP embeddings for each sample, assigned cluster identities, and quantified the proportion of cell types within each sample (**Figure 1D**). The relative proportion of each population varied when comparing Xenium and scRNA-seq data, which was most likely due to technical artifacts introduced during scRNA-seq. For example, scRNA-seq specimens used whole thickness tissue dissociation, which may skew cell proportions^57–61^. In contrast, Xenium used a fixed 5µm cross-section of each intestinal region imaged in a z-stack, such that cell proportions are not altered and are fixed in their near-native state but are dependent upon the plane of tissue observed.

Interestingly, proportions in scRNA-seq data showed slightly higher variation in the smooth muscle cell (SMC) and immune populations across developmental time (**Figure 1B** - dark green and pink) compared to Xenium, where these populations remained consistent over the period of developmental time observed (**Figure 1D**). Again, the difference across platforms may be attributed to technical differences. For example, the immune cells are more easily isolated in single-cell dissociations than other populations, which may explain why there was a higher proportion of immune cells identified in the scRNA-seq data. The epithelium had the largest difference in the number of cells identified between the two methods, with the Xenium data containing almost double the proportion of epithelial cells (scRNA-seq: 11.4-18.1%; Xenium: 30.3-36.5%) (**Figure 1B and D** - gold). In both methods the fibroblasts account for roughly half of the cells analyzed (scRNA-seq: 47.2-52.4%; Xenium: 37.9-49.3%; **Figure 1B and D** - navy).

Despite these differences, we observed that data from both technologies identified the muscularis mucosa between 80-105 DPC (**Figure 1B and D** - bright green), consistent with prior studies showing that the muscularis mucosa begins to differentiate below the crypt domain during this window of development^16,18,62^. Overall, the spatial data allowed visualization of the tissue architecture and composition that was obscured in previous dissociated scRNA-seq datasets.

### Identification of diverse fibroblast populations within the developing intestine

Using human fetal intestinal scRNA-seq^4,32–35^ (**Figure 2A, Supplemental Table 1**) we extracted the mesenchymal fibroblast populations^6,7^ (*VIM+, COL1A1+, COL1A2+, THY1+,* and *CDH2*+; **Supplemental Table 3**), and computationally removed four populations: 1) *ACTA2+/TAGLN+* smooth muscle cells that made up the inner circular and outer longitudinal muscle layers of the muscularis externa; 2) VSMCs/pericytes (*RGS5*+, *KCNL8+, ABCC9+, BGN^hi^, CCDC3^hi^*); 3) actively cycling cells (*MKI67*+, *TOP2A+, PCNA+*); and 4) Interstitial Cells of Cajal (ICCs) (*KIT+, ANO1+*) (**Supplemental Table 4**). After batch correction, the fibroblasts of the scRNA-seq data clustered broadly by developmental stage (**Figure 2A**), and within each developmental stage, we observed clustering into 3 major sub-populations: subepithelial cells (SECs), lamina propria fibroblasts (LPFs), and sub-mucosal fibroblasts (SMFs). Within the most developmentally advanced samples (110-142 DPC), we also observed a *CXCL13*+ fibroblast population (**Figure 2C**). The *CXCL13*+ population was rare, comprising only 5.5% of the cells in the scRNA-seq data. Although the Xenium data did not separate by age (**Figure 2B**), all major fibroblast sub-populations (SEC, LPF, and SMF) were identified, as were the rare *CXCL13*+ fibroblasts (**Figure 2B, 2D**). We identified highly expressed genes within these sub-populations in both scRNA-seq and Xenium data (**Figure 2E-F**), which include: SEC fibroblasts – *PDGFRA^hi^, F3;* LPF fibroblasts – *FABP5, ADAMDEC1, GPX3, ALDH1A3;* SMF fibroblasts – *SHISA3, FIBIN, C1QTNF3;* and *CXCL13*+ fibroblasts – *CXCL13*.

**Figure 2:**
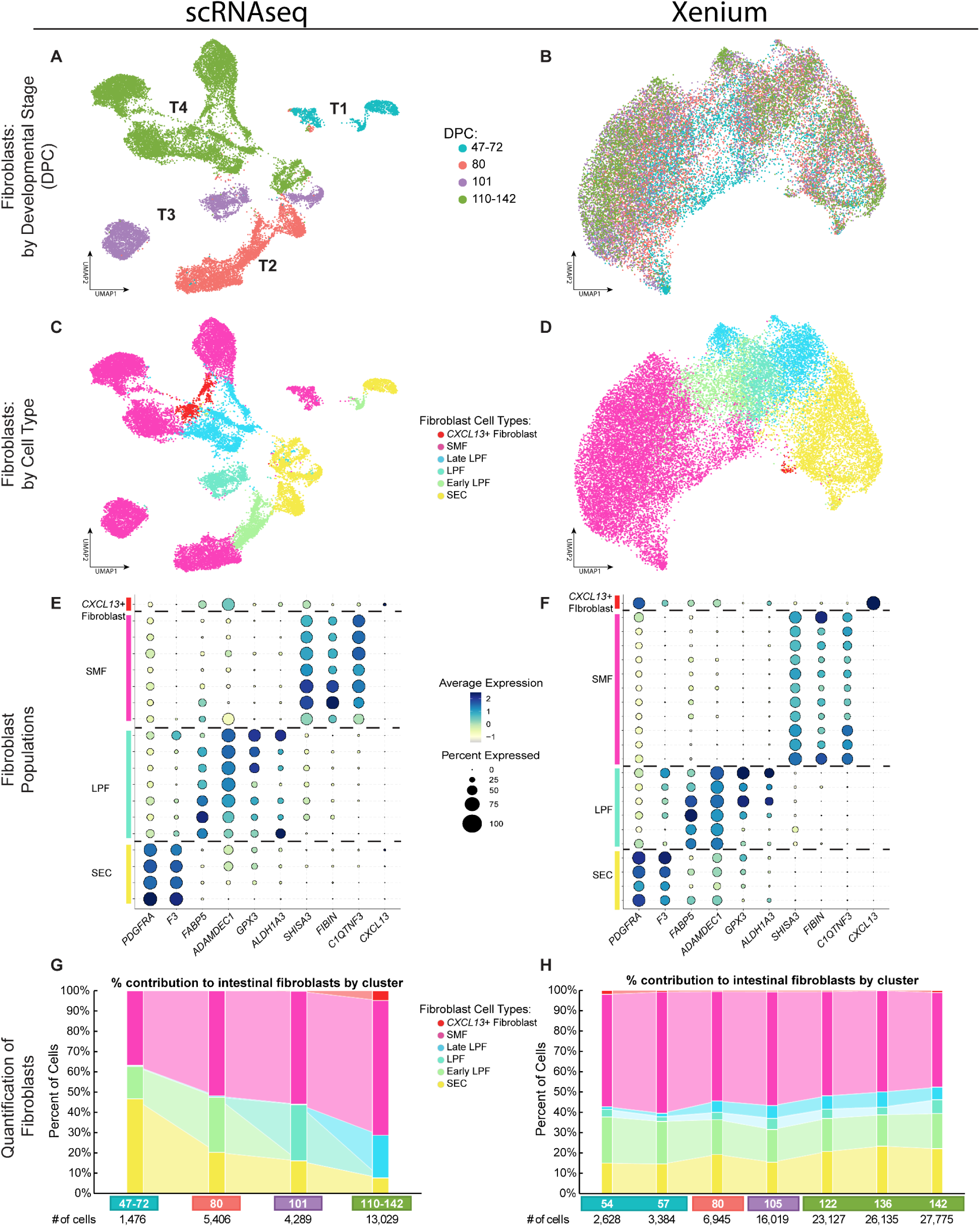
Identification of diverse fibroblast populations within the developing intestine using scRNA-seq and Xenium. UMAP projections summarizing the extracted fibroblast populations observed in human fetal small intestine as determined by **A)** scRNAseq and **B)** Xenium spatial transcriptomics, pseudo-colored by developmental stage. **C)** scRNAseq and **D)** Xenium UMAPs pseudo-colored colored by cell type. Dot plots displaying the markers used to annotate individual clusters for the **E)** scRNAseq and **F)** Xenium data. Stacked bar graphs describing the fibroblast cell population composition of the human fetal small intestine at various time points through development (47-142 DPC) for either the **G)** scRNAeq dataset or **H)** Xenium dataset. Sample labels below the graphs in **G-H** are colored to indicate the developmental stage (DPC) each sample falls within. The number of fibroblast cells included in each sample is indicated below the sample age. Cell populations are color matched in **C-H**; SECs (yellow), early LPFs (mint green), co-expressing LPFs (light teal), late LPFs (cyan), SMFs (pink), and *CXCL13*+ Fibroblasts (red).

We have previously described SECs in the developing human intestine, marked by high expression of *PDGFRA* and *F3*^1,5,7^. These cells are immediately adjacent to the epithelium, line the crypt-villus axis, and have been described in the literature as telocytes, telopodes, Interstitial Cells of Cajal Like Cells/ICLCs, fibrocytes, and subepithelial myofibroblasts^9,11,13,15,17,19,21,23,63^. Interestingly, in the scRNA-seq data, we observed changes in the proportion of fibroblast sub-populations as development progressed (**Figure 2G**), whereas the cell proportions were consistent across time in the Xenium data (**Figure 2H**), suggesting that dissociation might have altered cell proportions of fibroblasts when preparing cells for sequencing. The Xenium data was confirmed with fluorescent *in situ* hybridization (FISH) using independent biological specimens (**Supplemental Figure 2A, B**), where we observed that the *F3*+ SECs were present in all the FISH imaging as a single layer of cells adjacent to the epithelium in samples spanning the age range 57-125 DPC (**Supplemental Figure 2**).

Thus, we have used scRNA-seq and Xenium data to identify distinct mesenchymal cell populations that could be broadly divided into two categories: the smooth muscle cells (SMCs) consisting of the outer longitudinal/inner circular SMCs and the muscularis mucosa (**Figure 1**), and the non-muscle fibroblast populations which include SECs; LPFs; SMFs; and *CXCL13*+ fibroblasts (**Figure 2**). The data captured not only established intestinal mesenchymal populations, but also populations in the process of forming (i.e. muscularis mucosa), and they highlight the diverse nature of the intestinal mesenchyme and fibroblasts while providing an atlas of cellular distribution during development.

### Localization of SECs, LPFs, SMFs and *CXCL13*+ cells within the intestine

#### Localization of the SEC (F3+) population

The intestinal subepithelial cells (SECs) made up a single cell layer immediately adjacent to the intestinal epithelium and have been described in the human intestine as having high *PDGFRA, F3* and *DLL1* expression^1,5^, although we observed *DLL1* expression in both SECs and lamina propria fibroblasts (LPFs) fairly evenly within our Xenium data. SECs were identified at all developmental stages interrogated by Xenium (54-142 DPC) (**Figure 3B1-B4**), were confirmed to reside directly adjacent to the epithelium, and expressed *F3* robustly. We also found that, while *F3* appeared to be specific to the SEC population within the fibroblasts, *F3* was expressed at low levels in the epithelium at early time points (54 and 57 DPC), and in the region of the muscularis externa after 105 DPC (**Figure 3B1-B4**). Localization of the SECs observed within the Xenium dataset was confirmed using fluorescence in situ hybridization (FISH) to probe for *F3* mRNA expression in independent biological samples (57-125 DPC, **Supplemental Figure 2A-B** - yellow).

**Figure 3:**
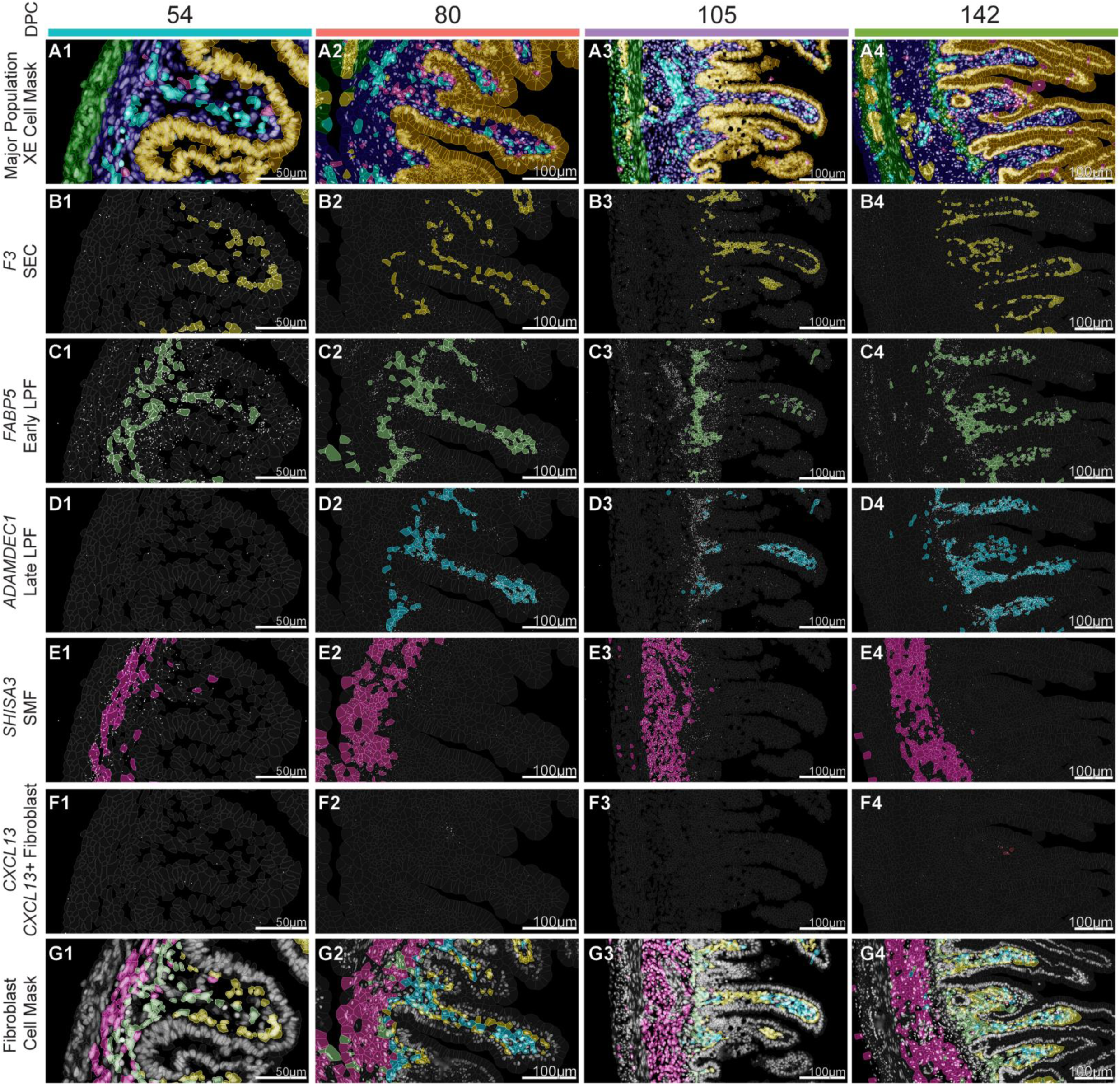
Localization of fibroblast sub-populations using Xenium spatial transcriptomics. Expression of proposed human fetal intestinal fibroblast markers over developmental time in the duodenum. **A)** Landmark populations of the muscularis externa and muscularis mucosa (green), epithelium (orange), fibroblasts (navy), endothelium (cyan), ENS (yellow), and immune cells (pink) are provided for localization reference. Fibroblast populations identified in the sequencing data include **B)** SEC (yellow), **C)** *FABP5+* early LPF (mint green), **D)** *ADAMDEC1+* LPF (cyan), **E)** *SHISA3* (pink), **F)** *CXCL13+* fibroblast (red). White dots indicate the expression of the indicated marker transcript. **G)** Composite mask displaying all fibroblast cell types. Scale bars = 50-100µm. Colored bars indicate the developmental stage (DPC) each sample falls within. DAPI depicted in grey in **A** and **G**.

#### Localization of the LPF (FABP5+/ADAMDEC1+) population

The lamina propria fibroblast (LPF) population was located within the villus core beneath the layer of SECs and spans the region from the villus shelf/crypt to the villus tip. During development, these cells expressed *FABP5, ADAMDEC1, ALDH1A3, and GPX3* (**Figure 2E-F**). We have previously reported that the LPF population adjacent to the SEC population expresses *GPX3* in tissue 132 DPC^1^, although, here we found that *GPX3* is a poor LPF marker in the early stages of development, where it appeared by 80 DPC but did not show consistent and robust expression until 122 DPC. However, *GPX3* serves as an excellent LPF marker in adult tissue, when other LPF markers *ADAMDEC1* and *FABP5* are not as highly expressed (**Supplemental Figure 3**).

Interestingly, during early development (47-57 DPC) we observe dynamic *FABP5* and *ADAMDEC1* expression, with robust *FABP5* expression (**Figure 3C1**) and little *ADAMDEC1* expression early (**Figure 3D1**). LPF cells begin to co-express *ADAMDEC1/FABP5* at later stages (from 80 DPC onward) (**Figure 3C2-C4** and **Figure 3D2-D4**), until expression of *FABP5* was largely lost and *ADAMDEC1* became the predominant marker. The combined *FABP5*+ and *ADAMDEC1*+ populations of LPFs account for 24.97-30.3% of the fibroblasts identified in the Xenium data during development (**Figure 2H**).

Using FISH, we observed widespread *FABP5* mRNA expression in an independent 57 DPC sample, and *ADAMDEC1* mRNA was absent. By 82 DPC, *ADAMDEC1* was expressed clearly and specifically in the LPFs, but not in *F3*+ cells (**Supplemental Figure 2A-B**). Within the fibroblast populations, *FAPB5* was highly specific to the LPF population; however, *FABP5* was also expressed in several other non-fibroblast cell types including pericytes (*RGS5*+), endothelium (*PECAM1*+), epithelium (*CDH1*+), and smooth muscle layers (*ACTA2*+), which accounted for a majority of *FABP5* expression from d82 onwards. The *FABP5*+ fibroblast population however is distinctly *FABP5+/PECAM1*-*/RGS5*-*/ACTA2*-*/CDH1*-, meaning it is distinct from any of the other *FABP5* expressing populations in the lamina propria. Thus, *FABP5* was not very useful as an imaging marker, but was sufficient to identify early LPFs when other populations can be filtered from the analysis, such as with sequencing data (**Supplemental Figure 2A-D**).

#### Localization of the SMF (SHISA3+) population

The sub-mucosal fibroblasts (SMF) reside between the crypts/muscularis mucosa and the outer muscular layers (muscularis externa) and could be identified by the markers *SHISA3, FIBIN,* and *C1QTNF3* (**Figure 2E-F**), although *SHISA3* was the most consistently robust marker across all developmental stages. Early in development, the SMFs populated a loosely formed layer that was several cells thick, while later in development (105 DPC onward) the SMF population expands proportionally to fill the entirety of the space between the muscularis mucosa and the inner circular muscle layer (**Figure 2F**, **Figure 3E1-E4**). FISH staining shows *SHISA3* expressing SMFs populating the submucosa, similar to the Xenium data (**Supplemental Figure 2C-D**). Transcriptionally similar cells have also been described as collagen and decorin (DCN) expressing previously in the sequencing data^1^.

#### Localization of the CXCL13+ population

The *CXCL13*+ fibroblast population is a small and rare population that was first observed after 101 DPC in the scRNA-seq data. Marked by expression of *CXCL13* in the sequencing data, the cells of this population are identified by *CXCL13* expression in the Xenium data and located near the crypt-base SECs, at the villus shelf, or mid-way up the villus immediately adjacent or integrated within the SEC layer (**Figure 3F4**). The FISH data confirmed the expression of *CXCL13* in rare cells of an independent 125 DPC sample (**Supplemental Figure 2E-F**).

Taken together, Xenium spatial imaging confirmed the localization of 4 non-smooth muscle populations of intestinal fibroblasts in the developing human intestine. This included the previously described SECs (*PDGRFA^HI^, F3+*) adjacent to the epithelium, the villus core-localized LPFs which we further described temporally based on marker distribution (*FABP5+, ADAMDEC1*+), the SMFs (*SHISA3*+) which are located in the submucosa between the muscularis externa and the crypt epithelium, and a rare *CXCL13*+ fibroblast that largely localized to the villus shelf/upper-crypt region of the mucosa.

## Creating a spatial atlas of pluripotent stem cell derived human intestinal organoids

After identifying and mapping broad cell classes (epithelium, fibroblasts, endothelial, ENS, and immune) (**Figure 1**) and a more detailed atlas of fibroblast sub-type gene expression and localization in the developing human intestine (**Figures 2-3**), we set out to use this information to interrogate the cell types and organization of pluripotent stem cell-derived human intestinal organoids (HIOs). We have previously used different methods to differentiate HIOs with varying levels of cellular complexity *in vitro.* For example, HIOs grown with EGF possess limited cellular complexity *in vitro* (containing immature epithelium and mesenchyme), whereas HIOs grown with EREG co-differentiate multiple lineages (epithelium, mesenchyme, endothelium, ENS)^55,64^.

When grown *in vitro,* HIOs possess limited organizational complexity; for example, they lack a crypt-villus axis. However, after transplanting under the kidney capsule of an immunocompromised mouse, HIOs gain both cellular and organizational complexity, develop a crypt-villus axis, and acquire distinct smooth muscle layers^63,64^. Therefore, we interrogated both EGF-and EREG-grown transplanted HIOs (tHIOs) using the Xenium human fetal small intestinal probe set (**Supplemental Table 2**).

Given that the human-specificity of the probeset resulted in transcript counts that were comparatively low in murine cells and had a decreased variety of transcripts, host mouse cells were easily identified in the tHIOs and removed from further analysis (**Supplemental Figure 4**). These cells were identified as distinct clusters using graph-based clustering due to the low feature numbers. Moreover, the majority of cells in the low expression cell clusters were located at the periphery of the HIOs, where murine tissue was still attached to the HIO following dissection and could be confirmed visually (**Supplemental Figure 4**). Excluding cells based on feature counts rather than using a region-of-interest (ROI) based method allowed us to preserve the fringe human cell populations that are ‘invading’ the mouse tissue for the dataset while also removing any mouse cells that ‘invaded’ the tHIOs (**Supplemental Figure 4F and G**).

Using methods described for **Figure 1**, the major cell classes (epithelium, fibroblasts, endothelium, ENS, and immune) were identified in tHIOs derived using two different HIO differentiation methods^55^ (**Figure 4**). In both EGF-grown and EREG-grown tHIOs, we identified all of the major cell classes, except human-derived immune cells, which are known to be absent from both kinds of tHIOs^45,55,65^ (**Figure 4**). EGF- and EREG-grown tHIOs also both possessed multiple smooth muscle layers including distinct muscularis externa and muscularis mucosa, as expected based on our previous studies^66^ (**Figure 4D and 4E**).

**Figure 4:**
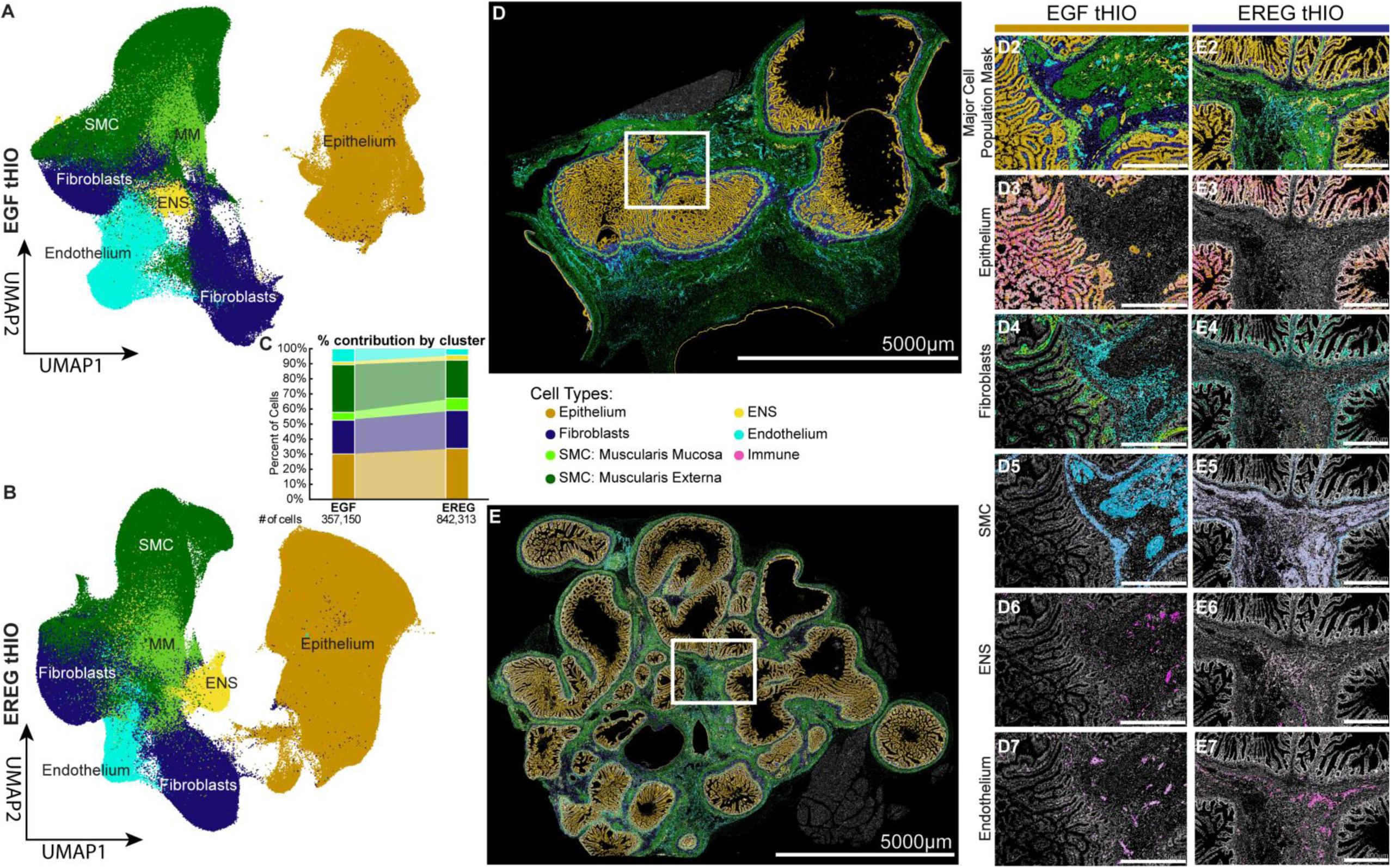
Identification and comparison of EGF-tHIO and EREG-tHIO major cell class composition. UMAP projections of the major cell class composition in a Xenium image of both an **A)** EGF-tHIO and **B)** EREG-tHIO. **C)** Stacked bar graph summarizing the percent cellular composition of the EGF-tHIO (left) and EREG-tHIO (right) samples. Cell ID masks generated in XE depicting **1)** the entire sample acquired for the **D)** EGF-tHIO and **E)** EREG-tHIO. Scale bar = 5000µm. The white box indicates the ROI depicted in **D2/E2-D7/E7. D2-7 and E2-7** display the cell sub-type distribution of each major cell class identified using the Xenium panel visualized as **2)** the major cell class mask, **3)** the epithelium sub-types, **4)** the fibroblast sub-types, **5)** the SMC-related sub-types, **6)** the ENS sub-types, and **7)** the endothelium sub-types. Scale bar = 500µm Cell classes are color matched in **A-E**; epithelium (gold), fibroblasts (navy), muscularis mucosa (MM, light green), SMC and pericytes (dark green), ENS (yellow), and endothelium (cyan). No immune cells were observed in either tHIO sample. DAPI staining of cell nuclei depicted in grey.

Interestingly, while the EGF and EREG HIOs showed significant differences in their average composition pre-transplant under the mouse kidney capsule^53–55^, the overall cellular composition and tissue architecture were similar after transplantation (**Figure 4C-E**). Our finding of a robust population of human-derived endothelial cells in the EGF-grown tHIO Xenium data was somewhat surprising given that prior data has shown that endothelial cells are often present in EGF-grown HIO cultures during the first ∼week *in vitro* but are lost over time^45^ and are generally missing from EGF-tHIOs^54^, with the endothelial populations within tHIOs often being characterized as infiltrating from the mouse instead of human-derived. On the other hand, given our past data showing that endothelial cell survival in EGF-grown HIOs can be enhanced with VEGF^45^, it is possible that the human-derived endothelial cell population is highly variable in EGF-grown HIOs.

With respect to the major cell classes, cell composition and architecture of the tHIOs was remarkably similar to the developing human small intestine when comparing the Xenium datasets. The tHIOs were composed of ∼30% epithelium, compared to ∼33.7% in the intestine data. Fibroblasts in the EGF-grown and EREG-grown tHIOs made up 22.4% and 25.2% of the samples, respectively, compared to an average 32.8% in the small intestine data. The tHIOs exhibited a higher proportion of SMCs (ME and MM combined) when grown in either EGF (37.1%) and EREG (33.4%) than the tissue (22.5%). The remaining populations - the ENS and endothelium - appeared to also have similar proportions across organoids and tissue. The EGF-grown and EREG-grown tHIOs had 2.0% and 3.4% of cells identified as the ENS and 8.7% and 4.2% identified as endothelium, respectively. This was similar to the Xenium small intestine data which identified 3.6% of cells as ENS and 7.7% as endothelium while the sequencing data had on average a slightly larger ENS component (5.5%) and a smaller endothelial component (3.4%) (compare **Figure 1B and D** and **Figure 4C**).

We observed that tHIOs self-organized into a multi-layered structure with organization similar to the developing human intestine. While each tHIO possessed multiple lumens, within the structure of each lumen-containing organoid unit there was a single epithelial layer almost entirely organized in a crypt-villus like axis complete with multiple functional domains including the ISC-containing crypts distinct from the absorptive populations of the villus, a lamina propria composed of fibroblasts, a layer several cells thick of muscularis mucosa, a thicker submucosal fibroblast layer that contains both the ENS and the endothelium, and two distinct muscularis externa-like layers throughout which more ENS and endothelium could also be found (**Figure 4D-E**).

## Identification of diverse fibroblast populations within tHIOs

We next examined the tHIO data to determine if SECs, LPFs, SMFs, and *CXCL13*+ fibroblasts were present and appropriately organized within tHIOs.

In general, all the fibroblast subtypes identified *in vivo* were also present in both EGF-grown and EREG-grown tHIOs, except for the late-developing *CXCL13*+ fibroblast population which were found in neither tHIO (**Figure 5**). Of note, the two conditions are very similar in their distribution of SECs, SMFs, and LPFs. LPFs account for 18.8% and 19.3% of the fibroblasts in EGF-grown and EREG-grown tHIOs respectively, with a trend of the EGF-grown tHIO towards the earlier/ more immature *FABP5*+*/ADAMDEC1-* population while the EREG-grown tHIOs appeared to have a higher proportion of the more mature *FABP5-/ADAMDEC1+* LPF population (**Figure 5C**). The two tHIO conditions have almost identical proportions of SECs (EGF: 24.2%, EREG: 24.1%) and similar SMF contributions (EGF: 42.0%, EREG: 47.8%) when corrected for all fibroblasts within each sample. There was also an ‘undefined’ population of cells in the tHIOs that expressed fibroblast markers *VIM*, *COL1A1*, *COL1A2*, but none of the markers for the more specific fibroblast cell types (**Figure 5A-C, navy**; EGF: 15.0%, EREG: 7.1%). This population was likely an artifact of the culture system, as these cells appeared to be scattered evenly ‘outside’ of the individual organoid structures in the spaces between the muscularis externa of the organoids and the edge of the transplanted HIOs rather than incorporated into the intestine-like architecture of the organoid units or displaying an organized structure or localization of their own.

**Figure 5:**
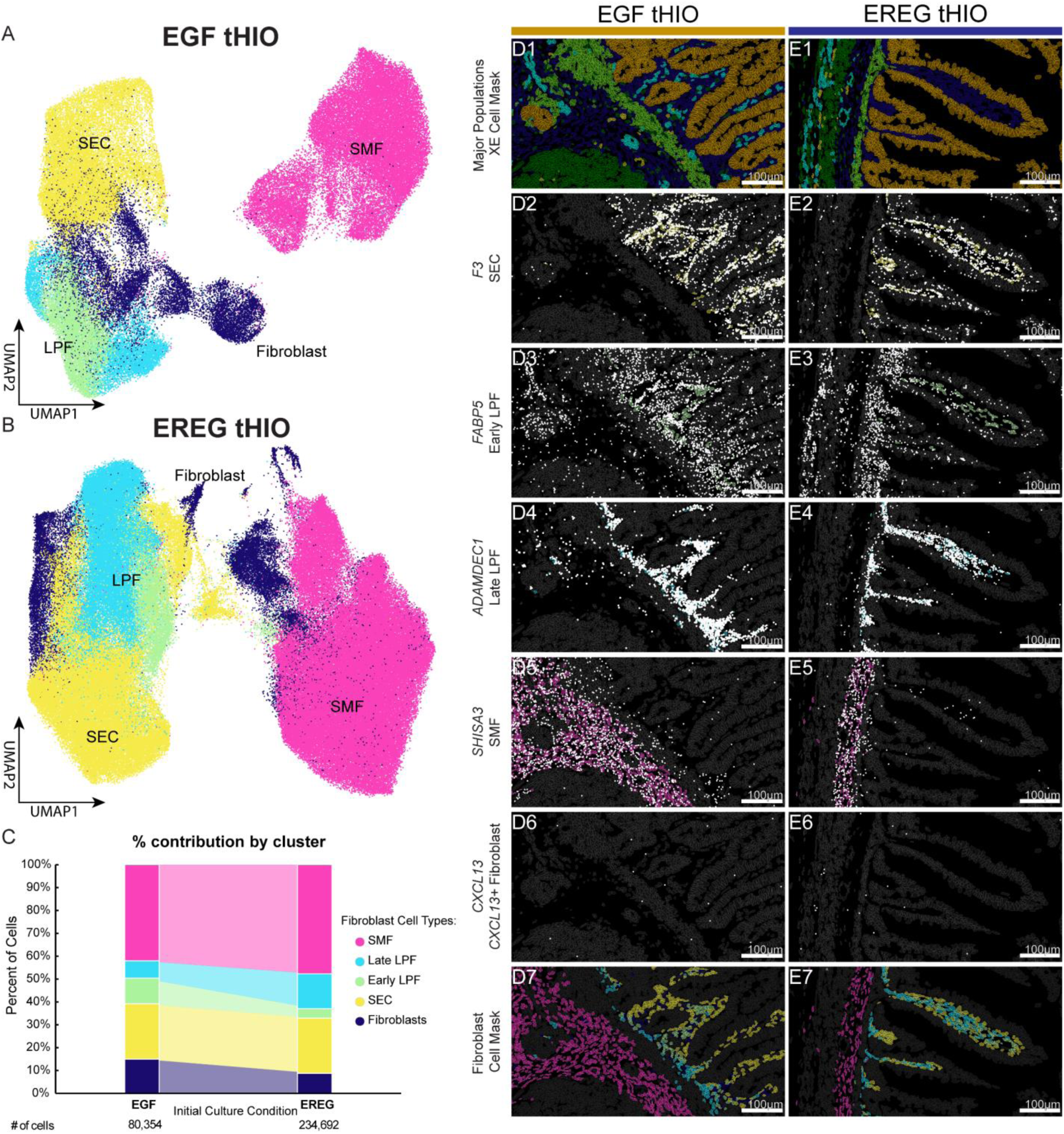
Fibroblast populations within tHIOs mimics distribution within human tissue. UMAP projections of the fibroblast cell class composition in a Xenium image of both an **A)** EGF-tHIO and **B)** EREG-tHIO. **C)** Stacked bar graph summarizing the percent cellular composition of the EGF-tHIO (left) and EREG-tHIO (right) samples. Cell ID masks generated in XE depicting **D)** EGF-tHIO and **E)** EREG-tHIO samples. **1)** XE cell mask summarizing the major cell classes for the tHIOs to landmark fibroblast population locations. Fibroblast sub-type distribution is depicted as **2)** the *F3*+ SECs, **3)** the *FABP5*+ early LPFs, **4)** the *ADAMDEC1*+ late LPFs, **5)** the *SHISA3*+ SMFs, and **6)** the *CXCL13*+ Fibroblasts. **7)** depicts the composite mask summarizing all fibroblast cell types in the tHIOs. Scale bars = 100µm

The distribution and organization of the fibroblast populations within organoid units was similar to in tissue as well (**Figure 5D-E**). The SECs line the epithelium and in areas where villi are thick enough the LPF population was found in the villus core. The SEC marker *F3* was also expressed in the intestinal epithelium of the more immature tissue specimens (54 DPC) but not in specimens 80 DPC and onward (**Figure 3B**). The EGF-grown tHIOs have clear epithelial expression of *F3* while the EREG-grown tHIOs do not have epithelial *F3* expression to the same extent, suggesting that the epithelium of EGF-grown tHIOs may be less mature than the EREG-grown tHIOs.

The LPFs extended to the muscularis mucosa layer, occupying the crypt shelf region as well as the villus core (**Figure 5D3-4, E3-4**). Despite markers of various cell types indicating relative immaturity, both EGF-and EREG-grown tHIOs have a distinct, well-developed muscularis mucosa within almost every organoid unit, despite the muscularis mucosa in the tissue developing much later (122 DPC, **Figure 1 and 3**).

The SMFs populated the region between the muscularis mucosa and the outer muscle layers, regardless of the distance between them, with varying thickness (**Figure 5D5, E5**).

The *CXCL13*+ fibroblasts were not observed in either tHIO model, and *CXCL13* expression overall was rare. Given the relatively immature state of tHIOs, the lack of the rare, late-onset population was not surprising. Despite the defined muscularis mucosa indicating a more mature state, the overall transcriptional identity of other populations that demonstrated temporal differences leans more immature, with the EGF-derived HIOs trending towards being more immature than the EREG-derived tHIOs.

Taken together, our data shows that except for the late-onset *CXCL13*+ fibroblast population, all of the fibroblast sub-populations identified in the scRNA-seq and Xenium data from the fetal intestine are present in EGF-grown and EREG-grown tHIOs and have a similar spatial organization when comparing *in vitro* differentiated organoids to *in vivo* tissue. This provides a framework for benchmarking organoids to *in vivo* tissue both in terms of cell heterogeneity and spatial organization, and it provides clues to how improvements can be made in organoid engineering.

## A spatial atlas of the adult duodenum

After characterizing the fibroblasts of the developing small intestine and using this to benchmark tHIOs, we examined an adult duodenum sample to determine if the fibroblast populations identified during development persisted into adulthood. We used the same probe set designed for the human fetal small intestine (**Supplemental Table 2**) to interrogate a full-thickness sample from the adult duodenum (44yo, female)^67^. We identified all major cell classes (**Figure 6A-E**) as well as the fibroblast subtypes identified in the fetal data (**Figure 6F-J**). At the level of individual genes, we noted some differences between fetal and adult; however, the identity and spatial localization of the fetal fibroblast populations were maintained in the adult.

**Figure 6:**
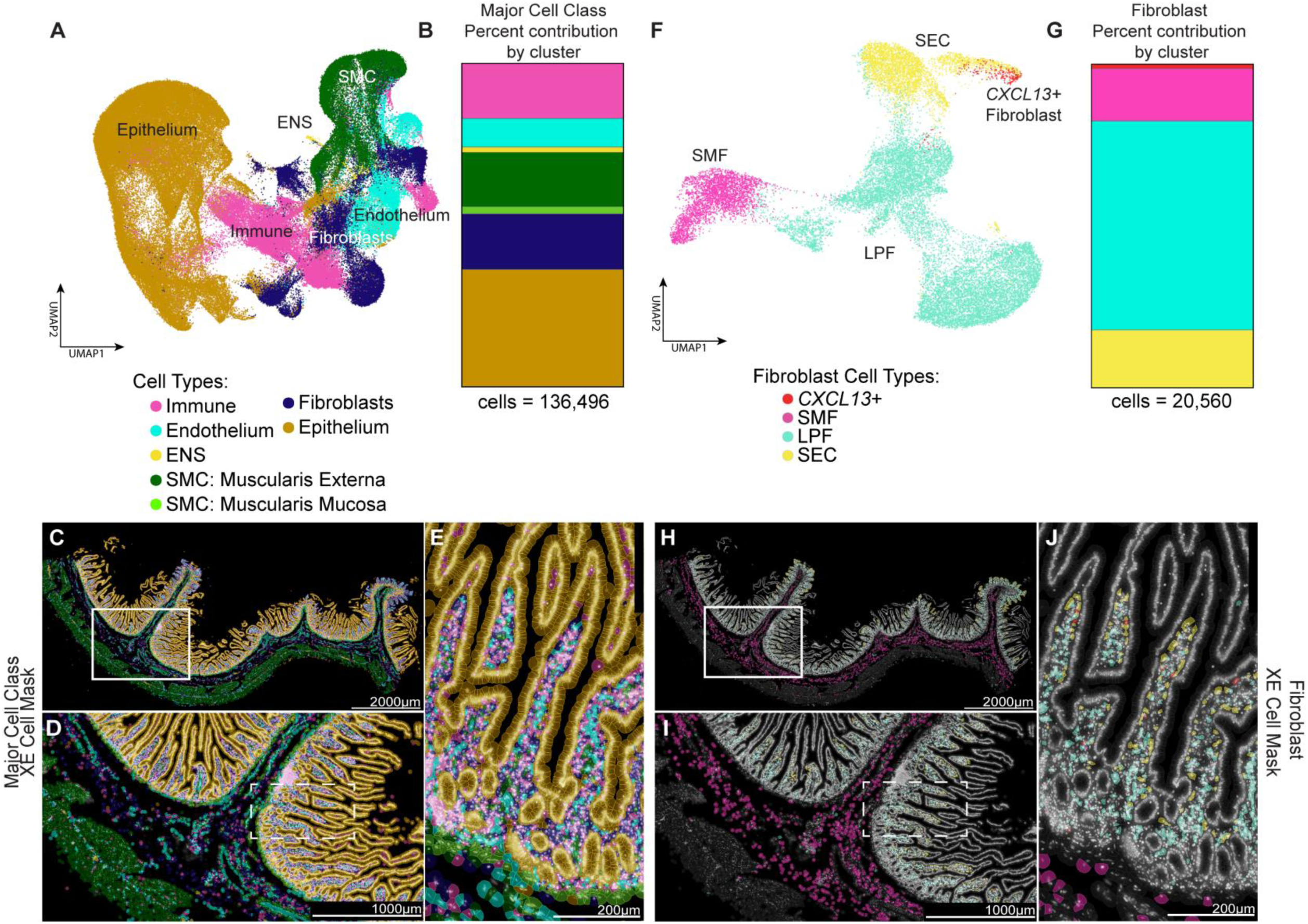
Summary of adult duodenum major cell class and fibroblast composition. UMAP projections of **A)** the major cell class composition and **F)** the fibroblast composition in a Xenium image of a full thickness cross-section of adult (44yo, female) duodenum. Stacked bar graphs summarizing the percent cellular composition of **B)** the major cell classes and **G)** the fibroblast cell types. Total cell number is included below the graphs. **C-E and H-J)** Cell ID masks generated in XE depicting **C)** major populations and **H)** fibroblasts for the entire sample acquired for the adult duodenum; scale bar = 2000µm. . **D and H)** Magnified ROI indicated in **C** and **H;** scalebar = 1000µm**. E and J)** Magnification of the crypt-villus axis to visualize the lamina propria composition; scale bar = 200µm. Cell classes are color matched in **A-E**; epithelium (gold), fibroblasts (navy), muscularis mucosa (MM, light green), SMCs (dark green), ENS (yellow), endothelium (cyan), and immune cells (pink). DAPI staining of cell nuclei depicted in grey. Fibroblast populations are color matched in **F-J**; SEC (yellow), LPF (cyan), SMF (pink), and *CXCL13+* fibroblasts (red).

We found that the full-thickness adult duodenum contained a similar proportion of epithelium (36.18%) to both the fetal (33.7%) and tHIO (30.1% and 33.8%) datasets and a smaller proportion (17.22%) of fibroblasts (fetal: 32.8%, EGF tHIO: 22.4%, EREG tHIO: 25.2%) (**Figure 6B**). Proportionally, the adult contained less muscularis mucosa (2.26%) and ENS (1.65%). There was about the same proportion of muscularis externa SMCs (16.80%) and endothelium (8.88%), and significantly more immune cells than observed in the fetal tissue (17.01%) (**Figure 6B**).

Perhaps the most significant differences observed between the fetal and adult Xenium data are within the fibroblast cell sub-types. We confirmed the presence of the four types of fibroblast identified in the fetal intestine and found that, while the SECs contribute a similar proportion of the fibroblast population in the fetal (16.4%) and adult (17.79%) and the *CXCL13*+ fibroblasts remain a relatively rare occurrence in the adult (1.33%), the LPFs made up the majority of fibroblasts in the adult tissue (64.67%), more than doubling the proportion of LPFs observed in the fetal tissue, which was relatively stable through development (25.2%, **Figure 6G** and **Figure 2H**). Correspondingly, the proportion of SMFs drops from being the major contributor in the fetal tissue (44.8%) to contributing only 16.20% in the adult. Furthermore, the spatial arrangement of the SMFs within the tissue during development was much more densely arranged than in the adult, where SMFs are more loosely scattered within the submucosa amid the extracellular matrix and large vessels (**Figure 6H-J**).

## Discussion

Understanding the cellular diversity within the intestine with spatial resolution using single cell approaches has recently shed light on rare and previously understudied populations in the murine and human intestine. In addition, fibroblasts – which are also referred to as stroma or mesenchyme – have historically been viewed as a broad population of cells that are important for structural integrity, and only recently have studies in mice shown that fibroblasts constitute a heterogeneous population with different functions^8,29–31,35,36,43,63,69–73^. Studies interrogating the human intestine have also revealed important cellular heterogeneity and localization of the intestinal fibroblasts^1–3,5,7^; however, the full complement of fibroblast cell types and their spatial complexity had yet to be mapped. Here, we leveraged scRNA-seq to design a custom probe set that targets the fetal human small intestine to capture the full diversity of cells using the Xenium platform. By profiling eight human specimens and two transplanted human intestinal organoid (tHIO) models, we provide—to our knowledge—the first tissue-scale, cellular-resolution atlas of human intestinal fibroblasts across a wide developmental continuum. This work reframes our understanding of fibroblast patterning during intestinal development, provides a practical framework for benchmarking stem-cell-derived intestinal tissues, and creates a resource for interrogating community-driven mechanisms of development.

## A cellular atlas of the human intestinal fibroblast lineages

During embryonic development, the mesoderm lineage gives rise to immune cells, endothelial cells, smooth muscle, and fibroblast/mesenchyme/stroma within the intestine. For the current work, we resolved four transcriptionally and spatially discrete fibroblast lineages--subepithelial cells (SECs), lamina propria fibroblasts (LPFs), sub-mucosal fibroblasts (SMFs), and rare *CXCL13+* fibroblasts (**Figure 7**). Each of these subsets occupied stereotypical, developmentally dynamic regions. Prior single-cell studies, including our own, have hinted at fibroblast heterogeneity and some positional information^1–3,5,7^, yet lacked sufficient resolution to create a map with stereotyped positions. Here, we demonstrate that (i) SECs form a continuous *PDGFRA*^hi^/*F3*^hi^ sheath immediately beneath the crypt–villus epithelium from as early as 7 weeks post-conception; (ii) LPFs (*FABP5+* and/or *ADAMDEC1+*) expand within the villus core and crypt shelf during villus elongation; (iii) SMFs (*SHISA3+*) progressively consolidate between the emerging muscularis mucosa and the muscularis externa; and (iv) *CXCL13+* fibroblasts arise late, clustering at the villus–crypt junction, an anatomical locus associated with lymphoid tissue formation.

**Figure 7:**
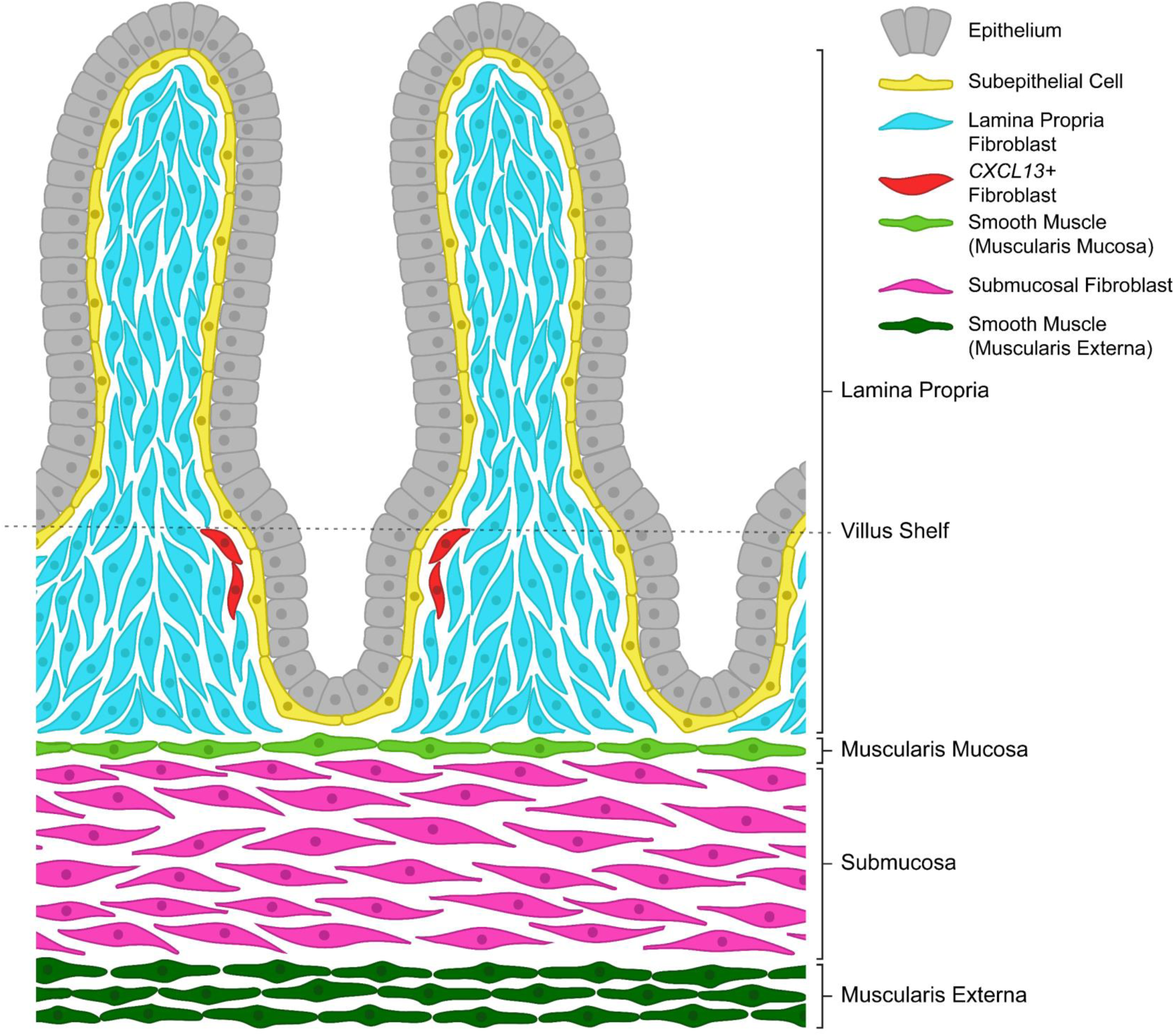
Schematic of human small intestinal fibroblast localization. Distribution of fibroblast populations identified by scRNAseq of human fetal small intestine. Fibroblast sub-types have distinct localizations: SEC (yellow) form a single cell layer adjacent to the epithelium, LPF (cyan) populate the remainder of the lamina propria, SMF (pink) populate the sub-mucosa, and *CXCL13+* fibroblasts (red) are found rarely within the SEC layer, usually at the crypt neck/villus shelf cupping a single crypt. Image includes epithelial cells (grey), SMC –muscularis mucosa (light green), and SMC – muscularis externa (dark green) for reference.

Indeed, *CXCL13+* cells have been identified previously using scRNA-seq^7^, the population increases in abundance though remains rare as development progresses, and are predicted to cross-talk with immune cells, suggesting a role in lymphoid tissue formation (Peyer’s Patches). Our findings also showed that within the lamina propria, LPFs can be subdivided into ‘early’ and ‘late’ since *FABP5* expression precedes that of *ADAMDEC1*, but then co-expresses with *ADAMDEC1.* Eventually, *FABP5* is largely absent from the LPFs, the population marked by expression of other genes *ADAMDEC1* and *GPX3*, suggesting *FABP5* marks an early or progenitor LPF state whose maturation warrants follow-up studies. This refined map reconciles several single cell studies that have identified diverse fibroblast populations and establishes molecular signatures and associated geographic locations within tissue compartments.

## Benchmarking and engineering intestinal organoids

Faithfully recapitulating human tissue heterogeneity and architecture has remained a key limitation of organoid technology. Using the Xenium spatial atlas data as a benchmark, we found that transplanted HIOs (tHIOs) generated under either EGF-or EREG-directed differentiation paradigms reproduce almost the full complement of fetal human fibroblast subtypes—including SEC, LPF, and SMF layers—with correctly ordered spatial topology. Subtle divergences were nevertheless evident: EGF-derived tHIOs retained epithelial *F3* expression, indicative of an immature epithelial state and exhibited a more immature *FABP5+/ADAMDEC1*-LPF profile, while EREG-grown tHIOs appeared to be more mature in both respects. On the other hand, both organoid models failed to generate *CXCL13+* fibroblasts. These findings underscore three principles.

First, cell-type abundance alone is an insufficient surrogate for fidelity; fibroblast positioning and stage-specific gene expression profiles should be considered. Second, the previous assumption that the presence of the more developed muscle layers—the muscularis externa and the muscularis mucosa—in the tHIO models to indicate a global maturation of all cell types present in the tHIOs does not seem to be accurate. Despite the general architecture of the tHIOs appearing to match more developed tissue morphology, the transcriptomic identities of some cell types appeared to remain relatively immature. Third, developmental immaturity and incomplete patterning remain as bottlenecks that need to be addressed in future experimental studies through modulation of growth-factor gradients, extracellular matrix composition, or co-culture with the major missing cell type in organoids, the immune cells. Work in this area is ongoing^66^. The atlas thus serves both as a quantitative benchmark for HIO quality control and as an instructive template for the next generation of tissue-engineered intestine.

## Limitations and future directions

Our study employs a 292-gene Xenium panel, inevitably biasing clustering towards pre-selected markers and limiting discovery of rare or novel states not identified prior to constructing the gene panel. Iterative panel expansion or complementary in-situ sequencing may reveal additional sub-specialization, such as telocyte heterogeneity along the villus axis or mesothelial–fibroblast intermediates. To this end, after developing the custom 292 gene panel used in this study, 10X Genomics developed an ‘off-the-shelf’ Xenium Prime 5K ultra-high-plex assay that explores 5,000 genes within a tissue. A limitation of this 5K panel is that specific or well characterized markers of individual cell types may be missing from the panel; however, it is likely that significantly enhancing the number of genes detected within a given cell will also enhance the fidelity of cell-type identification, regardless of whether specific genes are missing from the panel. Additionally, while our spatial atlas has now defined both cell types and locations within the fibroblast population, functionally interrogating these populations to define their role in intestinal development and homeostasis is a very important next step. Given that access to human tissue is limited, this further highlights the important roles for organoids as a model system to interrogate the human intestine. Finally, the fibroblast atlas, though comprehensive for the small intestine, must be extended to colon and to post-natal and adult stages to fully capture regional and temporal diversity.

## Conclusions

By uniting single-cell and spatial transcriptomics, we have refined the cellular cartography of intestinal cell types that underpin human intestinal development, and we have provided a quantitative scaffold for organoid engineering and disease modelling. Beyond cataloguing cell types, our work illustrates a generalizable strategy for translating transcriptional atlases into spatially resolved blueprints of human organogenesis. As spatial multi-omics technologies mature, such blueprints will be indispensable for deciphering cell-cell communication, predicting important cell types and locations for congenital and inflammatory disorders, and for engineering transplantable tissues that truly mimic human biology.

## STAR Methods

Key Resources Table

| REAGENT or RESOURCE | SOURCE | IDENTIFIER |
| --- | --- | --- |
| **Antibodies and Labels** |  |  |
| Rabbit anti-TAGLN /SM22 [1:500] | Abcam | ab14106; RRID: AB_443021 |
| Rabbit anti- αSmooth Muscle Actin (αSMA) [1:500] | Abcam | ab5694; RRID: AB_2223021 |
| Donkey anti-rabbit 488 [1:500] | Jackson ImmunoResearch | 711-545-152; RRID:AB_2313584 |
| NucSpot 750/780 [1:1000] | Biotium | 41038 |
| DAPI [1:1000] | Sigma | D9542 |
| **FISH Probes** |  |  |
| RNAscope Probe Hs-F3 | ACD | 407611 |
| RNAscope Probe Hs-ADAMDEC1-C2 | ACD | 478471-C2 |
| RNAscope Probe Hs-FABP5-C3 | ACD | 566111-C3 |
| RNAscope Probe Hs-SHISA3-C2 | ACD | 1314581-C2 |
| RNAscope Probe Hs-CXCL13-C3 | ACD | 311321-C3 |
| **FISH Fluorophores** |  |  |
| TSA Cy3 [1:1000] | Akoya | TS-000202 |
| TSA Cy5 [1:2000] | Akoya | TS-000203 |
| CF594 Tyramide [2μM] | Biotium | 92174 |
| **Biological Samples** |  |  |
| human fetal intestine | University of Washington Laboratory of Developmental Biology | N/A |
| human adult duodenum | University of California, San Francisco | N/A |
| **Critical Commercial Assays** |  |  |
| RNAscope® Multiplex Fluorescent Reagent Kit v2 | ACD | 323100 |
| Xenium Slides & Sample Prep Reagents | 10X Genomics | PN-1000460 |
| Xenium Cell Segmentation Staining Reagents | 10X Genomics | PN-1000661 |
| Xenium Standalone 300 Gene Panel | 10X Genomics | PN-1000563 |
| **Deposited Data** |  |  |
| Raw scRNA-seq data (human fetal duodenum) | Holloway et. al, 2020 | ArrayExpress: E-MTAB-9489 |
| Raw scRNA-seq data (human fetal jejunum) | Yu et. al, 2021 | ArrayExpress: E-MTAB-10187 |
| Raw scRNA-seq data (human fetal ileum) | Hung, et. al, 2019 | ArrayExpress: E-MTAB-9906, E-MTAB-10187, E-MTAB-9363 |
| Raw scRNA-seq data (human fetal duodenum, 70 DPC) | This paper | ArrayExpress: E-MTAB-15358 |
| Xenium – Human Intestinal Tissue fetal and adult (1/4) | This paper | 10.5281/zenodo.15838935 |
| Xenium – Fetal Human Intestine – EGF tHIO (2/4) | This paper | 10.5281/zenodo.15849955 |
| Xenium – Fetal Human Intestine – EREG tHIO (3/4) | This paper | 10.5281/zenodo.15850075 |
| **Software and Algorithms** |  |  |
| R Studio | Posit | varied<br>https://posit.co/downloads/ |
| R | CRAN | v4.1.2-4.4.1<br>https://cran.r-project.org/ |
| Seurat | Satija et al., 2015<sup>74</sup> | v4<sup>75</sup> and v5<sup>76</sup><br>https://satijalab.org/seurat/ |
| Cellranger | 10x Genomics | https://support.10xgenomics.com/single-cell-gene-expression/software/pipelines/latest/what-is-cell-ranger |
| UMAP | McInnes et al., 2018<sup>77</sup> | https://github.com/lmcinnes/umap |
| NIS Elements | Nikon | v5.24.01<br>https://www.microscope.healthcare.nikon.com/products/software/nis-elements |
| Xenium Explorer 2.0.0+ | 10X Genomics | https://www.10xgenomics.com/support/software/xenium-explorer/latest |
| FIJI | Schneider et al., 2012<sup>78</sup> | https://fiji.sc/ |
| Adobe Illustrator 2024 | Adobe | v28.1<br>https://www.adobe.com/ |
| Adobe Photoshop 2024 | Adobe | v25.3.1<br>https://www.adobe.com/ |
| Prism 10 | GraphPad | v10.4.1<br>https://www.graphpad.com/scientific-software/prism/ |
| Detailed methods and code for scRNA-seq and Xenium analysis | GitHub | [https://github.com/jason-spence-lab/Johnson et al 2025](https://github.com/jason-spence-lab/Johnson_et_al_2025) |
| R objects describing the Xenium data uploaded in the other three parts and associated scRNA-seq R objects (4/4) | Zenodo | 10.5281/zenodo.15850077 |
| **Other Chemicals and Reagents** |  |  |
| BSA | Millipore Sigma | A9647 |
| Histo-Clear II | National Diagnostics | HS-202 |
| UltraPure DNase/RNase-Free Distilled Water | ThermoFisher | 10977015 |
| Molecular Biology Grade Water | Corning | 46-000-CM |
| RNase Away | ThermoFisher | 700511 |
| FluorSave Reagent | Millipore Sigma | 345789 |

### Lead contact

Further information and requests for resources and reagents should be directed to and will be fulfilled by the Lead Contact, Jason R. Spence.

## Materials availability

All unique/stable reagents generated in this study are available from the Lead Contact with a complete Materials Transfer Agreement.

## Data code and availability

Sequencing data generated and used by this study is deposited at EMBL-EBI ArrayExpress. Datasets for human fetal intestine are detailed in **Supplemental Table 1 and STAR Methods** (ArrayExpress: E-MTAB-9489^1,5,45^, E-MATB-10187^5^, E-MATB-9906^5,46^, E-MATB-9363^45^ and E-MTAB-15358 - this paper).

Annotated Xenium datasets for the human fetal small intestine (Xenium – Human Intestinal Tissue fetal and adult (1/4): 10.5281/zenodo.15838935), EGF-derived tHIO (Xenium – Fetal Human Intestine – EGF tHIO (2/4): 10.5281/zenodo.15849955), EREG-derived tHIO (Xenium – Fetal Human Intestine – EREG tHIO (3/4): 10.5281/zenodo.15850075), and the processed R objects associated with the Xenium data (R objects describing the Xenium data uploaded in the other three parts and associated scRNA-seq R objects (4/4): 10.5281/zenodo.15850077) (**Supplemental Table 1**) can be found on Zenodo.

Code used to process both scRNA-seq and Xenium data can be found at https://github.com/jason-spence-lab/Johnson_et_al_2025.

## Experimental model and subject details

### Markers

The supervised lists of potential marker genes (**Supplemental Table 3** and **4**) were generated from the initial sequencing data (**Figure 1A** and **2A**) and mapped to at least one other publicly available human fetal intestine dataset^2,3^, then validated via FISH and Xenium imaging.

### Human subjects

Normal, de-identified human fetal intestinal tissue was obtained from the University of Washington Laboratory of Developmental Biology and normal, de-identified human adult duodenum tissue from research-consented deceased organ donors was obtained from the University of California, San Francisco (UCSF). Tissue from UCSF was obtained through Donor Network West in collaboration with the UCSF Viable Tissue Acquisition Lab (VITAL) Core as described previously^67^. All human tissue used in this work was de-identified and was conducted with approval from the University of Michigan IRB.

### Primary tissue collection, fixation and paraffin processing

Human intestine tissue samples were collected as ∼0.5 cm fragments and fixed for 24 hours at room temperature in 10% Neutral Buffered Formalin (NBF), and washed with UltraPure Distilled Water (Invitrogen, 10977-015) for 3 changes for a total of 2 hours. Tissue was dehydrated by an alcohol series diluted in UltraPure Distilled Water (Invitrogen, 10977-015). Tissue was incubated for 60 minutes each solution: 25% Methanol, 50% Methanol, 75% Methanol, 100% Methanol. Tissue was stored long-term in 100% Methanol at 4C. Prior to paraffin embedding, tissue was equilibrated in 100% Ethanol for an hour, and then 70% Ethanol. Tissue was processed into paraffin blocks in an automated tissue processor (Leica ASP300) with 1-hour changes overnight.

## scRNA-seq data analysis

The repositories of the raw scRNA-seq data of developing human fetal small intestine, duodenum, jejunum and ileum can be found in the STARl1lMethods Deposited Data section. Data matrices for further analysis were generated using the CellRanger pipeline under standard parameters by the University of Michigan Advanced Genomics Sequencing Core.

### Single cell library preparation and transcriptome alignment^1^

The single-cell RNA-seq sample library was prepared as previously described, with the 10x Chromium Controller using the v2 chemistry. Sequencing was performed on an Illumina HiSeq 4000 with targeted depth of 100,000 reads per cell. Default alignment parameters were used to align reads to the pre-prepared hg19 human reference genome provided by the 10X Cell Ranger pipeline. Initial cell demultiplexing and gene quantification were also performed with the default 10x Cell Ranger pipeline.

### Preprocessing/QC Filtering (Full Data)

Filtered gene expression matrices were imported into Seurat Version 4.1.0 in RStudio v1.4 running on R 4.1.2 for further analysis.

Quality control was conducted after merging 16 single Seurat objects. Cells were excluded if they contained less than 500 or greater than 8000 features, more than 10% mitochondria RNA, and the RNA reads were less than 500 or greater than 60000 to avoid the impact of outliers. The merged dataset was then normalized with a scale factor of 10,000 using the NormalizeData function. Highly variable features were identified using the FindVariableFeatures function. A Fast version of the Mutual Nearest Neighbors (MNN) was calculated to correct for batch effects using RunFastMNN. Uniform manifold approximation and projection (UMAP) was then constructed by RunUMAP function with mnn reduction created by RunFastMNN with the first 30 dimensions. Nearest neighbors were found by FindNeighbors function using MNN reduction. Cell clusters were identified by the original Louvain algorithm embedded in FindClusters with a resolution of 0.3.

### Cell Type Annotation (Full data)

The major classes of cell types in the human intestine [epithelium, fibroblast, immune, endothelium, enteric nervous system (ENS), and smooth muscles cells (SMC)] were manually annotated using major cell population markers listed in **Supplemental Table 3.** A cluster of contaminating red blood cells was identified and removed based on DEG analysis. Cell counts of each major cell type across time were calculated by categorizing sample by age within developmental stage (47−75 DPC; 80 DPC; 101 DPC; 110−135 DPC).

### Cell Type Annotation (Fibroblast subset)

The fibroblast subset of the scRNA-seq dataset that was analyzed for novel cell sub-types was defined after several rounds of quality control using Seurat version 5.2.1 in RStudio 2024.12.1. with R (version 4.4.1). First, the previously defined fibroblast population (**Figure 1A, Supplemental Table 3**) was extracted, and RNA assay was normalized using the SCTransform function with the percentage of mitochondrial reads regressing out in a second non-regularized linear, followed by the standard processing pipeline including principal component analysis (PCA), UMAP calculation, Nearest Neighbor discovery and cluster determination.

Then, according to the expression of gene markers listed in **Supplemental Table 4** and differential gene expression analysis identified by Wilcoxon Rank Sum test, we filtered out clusters that interfered with discovery of fibroblast cell types. Specifically, we removed the active cycling cells (*MKi67, TOP2A,* and *PCNA*) the smooth muscle cells (SMCs; *ACTA2* and *TAGLN),* pericytes (*RGS5, KCNJ8, KCNL8, ABCC9, BGN,* and *CCDC3)* and ICCs (*KIT* and *ANO1*). We removed these populations for two reasons. First, the clusters were significantly different from the remaining cells when viewed on a 3D-diffusion map, they caused all other clusters to clump together. Second, these populations (other than the cycling cells) are differentiated, non-fibroblast cell types by the transcriptomic definitions we established for this study. It took several rounds of QC to entirely remove each population, and for each round of our cell identification and quality control, we followed a similar pipeline and inspected gene expression using dot plots and feature plots. Cell counts (**Figure 2G**) of each sub cell type were calculated by categorizing samples by developmental stage (47−75 DPC; 80 DPC; 101 DPC, and 110−135 DPC).

## Xenium slide preparation and workflow

Xenium is a spatial transcriptomics method that permits imaging of hundreds-to-thousands of genes on a single FFPE sample at sub-cellular resolution (**Supplemental Figure 1A**). A custom fetal human small intestine Xenium probe panel was designed (**Supplemental Table 2**) based on conventional markers from the adult human, conventional markers from the mouse, and previously acquired or published sequencing data in the fetal human small intestine^1,3,4,33,34^.

As previously described^51^, archival tissue and tHIO FFPE samples were first prepared by first trimming excess paraffin from the paraffin blocks, which were then submerged in an ice bath for at least 30 minutes prior to sectioning. To prepare for section collection, the slide warmer was preheated 42°C and used to pre-warm positive-charged microscope slides topped with molecular biology grade water (Corning, Cat# 46-000-CM). The blocks were then sectioned at a thickness of 5μm, and serial sections were collected on the prepared slides. After allowing sections to flatten out, the water was removed, and sections were examined under the 4X and 10X objectives of a biological microscope to identify the optimal regions of the samples.

Using a single edge razor blade, the corresponding area of the paraffin block was lightly scored to obtain tissue sections small enough to fit into the 10.45mm by 22.45mm sample placement area of the Xenium slide. Xenium slides were removed from -20°C storage and allowed to come to room temperature 30 minutes prior to undergoing the same preparation as the microscope slides. Once the selected sections were placed on the Xenium slide, they were examined again to ensure that the fiducial frame was not obscured by any paraffin or tissue. Following water removal, sections were allowed to dry on the slide warmer for 3 hours before slides were moved to a microscope slide jar for transportation to the University of Michigan Advanced Genomics Core for processing and image acquisition using the Xenium Analyzer (10X Genomics, PN-1000481).

Due to the application of emerging technologies, the two different batches were processed for segmentation differently. In addition to the human fetal small intestine probe set, a segmentation kit (PN-1000661) was included to define cell boundaries of the second batch of samples which resulted in more accurate cell segmentation and as a result more accurate clustering both by the Xenium Explorer program and externally. Cell ids and transcript counts for the small intestinal samples were exported from Xenium Explorer (v2.0.0 and later) and analyzed using the pipeline described below. Images and digital cell masks were generated and exported using Xenium Explorer. Xenium image brightness (+100%) and contrast (+20%) were enhanced equally across all exported Xenium images in Adobe Photoshop CC 2024 before figure assembly in Adobe Illustrator CC 2024.

## Xenium data analysis pipeline

### Extraction of Xenium data for analysis

Xenium spatial transcriptomic data was exported from Xenium Explorer version (v2.0.0 and later). The cell-ids of the fetal small intestine (duodenum-jejunum-ileum) only were included by using the XE ROI selection tool to define the small intestinal segments and export only cells from those regions. The colon and (for 122-142 DPC samples) distal ileum were imaged but not analyzed in this study. The entirety of each tHIO sample, consisting of multiple lumen-containing organoid units were exported and analyzed as single samples per condition. The whole-thickness human adult duodenum sample was exported in its entirety and analyzed.

### QC of tHIOs: identification and removal of mouse cells

Before further analysis of the tHIOs could occur, the residual mouse cells were identified and removed from further analysis based on the variety of expressed genes and total probe counts per cell (**Supplemental Figure 3**). After validation of the removal of mouse cells and debris, the same pipeline was applied as for the human tissue samples.

### Xenium Cell Type Annotation (Full Data)

All Xenium data was loaded into R version 4.4.1 using a standard Seurat 5.2.1 pipeline. After loading individual sample data in, cells with 0 counts were filtered out and excluded from further analysis. Each individual sample was normalized using the SCTransform function which created a new assay with normalized transcriptomic expression values. Then PCA was performed on the top 30 PCs and was applied to UMAP construction. Nearest neighbors were found with the FindNeighbors function and cells were clustered by original Louvain algorithm at a 0.3 resolution using function FindClusters. Individual samples were carefully examined using dot plots and feature plots. Top 50 gene markers were generated by Wilcoxon Rank Sum test for each cluster for cell type annotation. The seven individual Xenium-sourced objects were then merged and cells were re-clustered following the same pipeline.

When generating UMAP visualizations, cells from the later two stages (105 DPC) and (combined 122, 136, and 142 DPC) were randomly down-sampled to 20,000 to reduce the computational burden (**Figure 1C**). Meanwhile, the whole dataset was analyzed and quantified to generate cell count data in each cell type across different ages for comparison. Major populations including epithelium, fibroblast, endothelium, ENS, SMCs, and immune cells were defined using the gene list in **Supplementary Table 3.**

Cell-IDs annotated using Seurat were imported back into Xenium Explorer and used to visualize the location of intestinal cell populations (105 DPC, **Figure 1F**). Each major cell class (epithelium, fibroblasts, SMCs, ENS, endothelium, immune) was then individually labeled to visualize sub-populations (**Figure 1G**). In order to carry out the labeling shown in **Figure 1G**, each population highlighted was computationally extracted, sub-clustered, and each cluster was annotated in Seurat, after which cell-IDs were re-imported into Xenium Explorer for visualization.

### Xenium Fibroblast Extraction

Different from extracting fibroblast population from a merged count matrices as we did for the single-cell sequencing data, we individually defined the fibroblast population for each sample after rigorous screening, before removing the proliferative cells, ICCs, SMCs, and pericytes from the fibroblast data based on marker gene expression (**Supplemental Table 4**). Then the individual fibroblast objects were merged, clustered, annotated, and quantified. Similar to the major cell type population Xenium data, the full fibroblast object was down-sampled to reduce computational burden for visualization of the UMAP and to prevent visual bias due to over-populating the object with the samples of developmental stages 122-142 DPC.

Stacked bar graphs depicting cell count distributions for major cell populations or fibroblast sub-types were generated in either Microsoft Excel (**Figures 1, 2, 4, and 5**) or GraphPad Prism (**Figure 6**). The annotated datasets were then re-loaded back into Xenium Explorer for visualization of the tissue architecture.

## Multiplex Fluorescent In Situ Hybridization (FISH) and Immunofluorescence (IF)

A list of probes, antibodies, fluorophores, and concentrations can be found in the Key Resources Table.

Archival tissues were selected from biologically distinct samples of similar age to at least one Xenium sample per developmental stage defined by the mesenchymal sequencing data. Paraffin blocks were sectioned to generate 5µm-thick sections within the week prior to performing *in situ* hybridization. All materials, including the microtome and blade, were sprayed with RNase-away solution prior to use. Slides were baked for 1 hour in a 60°C dry oven prior to deparaffinization and rehydration.

Tissue slides were deparaffinized in Histo-Clear II (National Diagnostics Cat#HS-202) twice for 5 minutes each, followed by rehydration in two washes of fresh 100% ethanol. The fluorescent in situ hybridization (FISH) protocol was then performed according to the manufacturer’s instructions (ACD; RNAscope multiplex fluorescent manual protocol, 323100-USM) with a 10-minute hydrogen peroxide step, a15-minute antigen retrieval step, and 30-minute protease plus treatment.

Immediately following the HRP blocking for the last channel of the FISH, slides were rinsed twice with PBS to remove excess detergent and then washed three times for 5 minutes in PBS, then transferred to blocking solution (5% Bovine Serum Albumin in PBS) for 1 hour at room temperature in a dark, humid chamber. Slides were then incubated with primary antibody (rabbit anti-TAGLN/SM22, ab14106 [1:500] or rabbit anti-αSMA, ab5694 [1:500]) suspended in blocking solution overnight at 4C in a dark, humid chamber. The following day, excess primary antibody was rinsed off twice with PBS before undergoing a series of 5 PBS washes of ten minutes each. Secondary antibody and nuclear dyes (donkey anti-rabbit 488 [1:500]), NucSpot 750/780 (1X), and DAPI [1:1000] were added to blocking solution and slides were incubated in a dark humid chamber at room temperature for 1 hour. Excess secondary antibody was rinsed off twice with PBS before going through another series of 3 PBS washes of 5 minutes each. Slides were mounted in FluorSave Mounting media (Millipore, 345789-20ML) and allowed to dry in the dark before being sealed. Once the seal was fully dried, slides were stored in the dark at 4C and imaged within a week.

All imaging was done using a Nikon AXR confocal after slides were brought to room temperature. Acquisition and the alignment of multi-track FISH images was performed using DAPI and NucSpot 750/780 (Biotium, 41038) in NIS Elements v5.24.01. A minimum of 3 ROIs per sample region were imaged. Images were acquired using a 20x air objective with 3.00x digital zoom as multipoint, multi-channel z-stacks using resonant scanning at 2048 x 2048, with fluorescence averaged over 4 images, and enhanced with Nikon Elements denoise.ai to further reduce background.

The z-stack series were then compiled into maximum intensity projections using FIJI^78^. For publication, a representative image from each sample was selected and the brightness was further enhanced in FIJI. Imaging parameters were kept consistent for all images within the same experiment and any post-imaging manipulations were performed equally on all images from a single experiment. Final figures were assembled using Adobe Illustrator CC 2024.

## Transplanted human intestinal organoids for Xenium imaging^55^

### Stem cell maintenance and differentiation

As previously published, stem cell lines were maintained in an undifferentiated state via media condition and regular splitting until differentiation. Stem cells underwent directed differentiation into definitive endoderm over a 3-day treatment. After three days, endoderm monolayers were differentiated into an intestinal identity. On days 4-6 of hindgut differentiation, spheroids budded from the monolayer and were collected. These spheroids were embedded in Matrigel as previously described^79^.

Organoid basal growth media was supplemented with epidermal growth factor (EGF) (100 ng/mL R&D Systems Cat#236-EG-01M) or Epiregulin (EREG) (10 ng/mL R&D Systems Cat#1195-EP-025/CF) with Noggin-Fc (100ng/mL, purified from conditioned media), and R-Spondin1 (5% conditioned medium) for the first three days of culture to pattern a proximal small intestine. On the third day after embedding, media was changed to basal growth media supplemented with EGF or EREG only (no additional Noggin or R-Spondin1) and remained in this media for the duration of the experiments with media changes every 5 days. Organoids were not passaged to avoid disrupting the development and spatial organization of the key cell types seen in EREG-grown HIOs. HIOs were cultured in vitro for at least 28 days then collected for transplantation.

### Mouse kidney capsule transplantation

The University of Michigan Institutional Animal Care and Use Committees approved all animal research. HIOs were implanted under the kidney capsules of immunocompromised NOD-scid IL2Rg-null (NSG) mice ^52–55^ (Jackson Laboratory strain no. 0005557). Between 1 and 3 HIOs were then surgically implanted beneath mouse kidney capsules using forceps. Mice were administered a dose of analgesic carprofen during the surgery and an additional dose after 24 hours. All mice were monitored daily for 10 days and then weekly until they were euthanized for retrieval of transplanted HIOs after 10 weeks.

### tHIO processing for staining and histology

The tHIOs were placed in 10% Neutral Buffered Formalin (NBF) for 24 hours at room temperature (RT) on a rocker for fixation. Fixed specimens were then washed 3x in UltraPure DNase/RNase-Free Distilled Water (Thermo Fisher Cat #10977015) for 30-60 minutes per wash depending on its size. Next, tissue was dehydrated through a methanol series diluted in UltraPure DNase/RNase-Free Distilled Water for 30-60 minutes per solution: 25% MeOH, 50% MeOH, 75% MeOH, 100% MeOH. Tissue was either immediately processed for paraffin embedding or stored in 100% MeOH at 4°C for future paraffin processing or whole mount staining. For paraffin processing, dehydrated tissue was placed in 100% EtOH, followed by 70% EtOH, and perfused with paraffin using an automated tissue processor (Leica ASP300) with 1 hour solution changes overnight. Tissue was then placed into tissue cassettes and base molds for sectioning. Prior to sectioning, the microtome and slides were sprayed with RNase Away (ThermoFisher Cat#700511).

5µm-thick sections were cut from paraffin blocks onto Xenium slides under RNAse-free conditions (10X Genomics, PN-1000460) and submitted to the University of Michigan Advanced Genomics Core for processing and image acquisition using the Xenium Analyzer (10X Genomics, PN-1000481). In addition to the human fetal small intestine probe set, a segmentation kit (PN-1000661) was included to define cell boundaries of the second batch of samples which resulted in more accurate cell segmentation and as a result more accurate clustering both by the Xenium Explorer program and externally.

## Human ethics approval

IRB: HUM00093465

## Acknowledgments

Research reported in this publication was supported by the National Cancer Institutes of Health under Award Number P30CA046592 by the use of the following Cancer Center Shared Resource: Single Cell and Spatial Analysis Shared Resource.

## Author Contributions

KJ and JRS conceived the study. KJ designed the Xenium panel. YT, and AW, KDW and KJ prepared tissue and Xenium slides and performed validation studies. KJ and XD analyzed and interpreted the data. XD performed all data curation and software management and performed bioinformatics analyses. RZ, OK, and IG provided the human tissue specimens used in this study. SGC assisted with figure design. JRS supervised and obtained funding for the research. KJ and JRS wrote the manuscript and XD wrote methods. All authors edited, read, and approved the manuscript.

## Abbreviations

scRNA-seq, FISH, IF, SEC, FFPE, Xenium, spatial transcriptomics

