## Supplemental Tables 1-4 for "Mapping mesenchymal diversity in the developing human intestine and organoids"

**Supplemental Table 1: Sample Demographics**

| **Technique** | **Sample Age** | **Sample Regions** | **Biological Sex** | **Data Publication** | **Data Availability** |
| --- | --- | --- | --- | --- | --- |
| scRNAseq | 47 DPC | Small Intestine | M | Holloway et. al, 2020 | ArrayExpress: E-MTAB-9489 |
| scRNAseq | 59 DPC | Small Intestine | F | Holloway et. al, 2020 | ArrayExpress: E-MTAB-9489 |
| scRNAseq | 70 DPC | Duo | M | This paper | ArrayExpress: E-MTAB-15358 |
| scRNAseq | 72 DPC | Duo | F | Holloway et. al, 2020 | ArrayExpress: E-MTAB-9489 |
| scRNAseq | 80 DPC | Duo-Jej-Ile | M | Holloway et. al, 2020;  Yu et. al, 2021;  Hung, et. al, 2019 | ArrayExpress: E-MTAB-9489, E-MTAB-10187, E-MTAB-9906 |
| scRNAseq | 101 DPC | Duo-Ile | M | Holloway et. al, 2020;  Yu et. al, 2021;  Hung, et. al, 2019 | ArrayExpress: E-MTAB-9489, E-MTAB-10187, E-MTAB-9906, E-MTAB-9363 |
| scRNAseq | 122 DPC | Duo | F | Holloway et. al, 2020 | ArrayExpress: E-MTAB-9489 |
| scRNAseq | 127 DPC | Duo | F | Holloway et. al, 2020 | ArrayExpress: E-MTAB-9489 |
| scRNAseq | 132 DPC | Duo-Ile | F | Holloway et. al, 2020;  Hung, et. al, 2019 | ArrayExpress: E-MTAB-9489, E-MTAB-9906, E-MTAB-9363 |
| Xenium | 54 DPC | Duo-Jej-Ile-Col | M | This paper | Zenodo: 10.5281/zenodo.15838935 |
| Xenium | 57 DPC | Duo-Jej-Ile-Col | M | This paper | Zenodo: 10.5281/zenodo.15838935 |
| Xenium | 80 DPC | Duo-Jej-Ile-Col | F | This paper | Zenodo: 10.5281/zenodo.15838935 |
| Xenium | 105 DPC | Duo-Jej-Ile-Col | F | This paper | Zenodo: 10.5281/zenodo.15838935 |
| Xenium | 122 DPC | Duo-Jej-Ile-Dist Ile-Col | F | This paper | Zenodo: 10.5281/zenodo.15838935 |
| Xenium | 136 DPC | Duo-Jej-Ile-Dist Ile-Col | M | This paper | Zenodo: 10.5281/zenodo.15838935 |
| Xenium | 142 DPC | Duo-Jej-Ile-Dist Ile-Col | F | This paper | Zenodo: 10.5281/zenodo.15838935 |
| Xenium | 10wk EGF iPSC-derived HIO | 100 ng/mL EGF  iPSC 72.3 | M | This paper | Zenodo: 10.5281/zenodo.15849955 |
| Xenium | 10wk EREG iPSC-derived tHIO | 10ng/mL EREG  iPSC 72.3 | M | This paper | Zenodo: 10.5281/zenodo.15850075 |
| Xenium | 44yo | Duo | F | This paper | Zenodo: 10.5281/zenodo.15838935 |
| FISH | 57 DPC | Duo-Jej-Ile-Col | F | N/A | N/A |
| FISH | 82 DPC | Duo-Jej-Ile-Col | F | N/A | N/A |
| FISH | 101 DPC | Duo-Jej-Ile-Col | M | N/A | N/A |
| FISH | 125 DPC | Duo-Jej-Ile-Col | M | N/A | N/A |

**Supplemental Table 2: 10X Xenium human fetal small intestine panel**

| **Gene** | ***Annotation*** | **Ensembl ID** |
| --- | --- | --- |
| ACTA2 | *Myofibroblasts / Myo-like* | ENSG00000107796 |
| ACTC1 | *OLM and cells between muscle layers* | ENSG00000159251 |
| ACTG2 | *Myofibroblasts / Myo-like* | ENSG00000163017 |
| ACVR1 | *RSTK Receptor* | ENSG00000115170 |
| ACVR1B | *RSTK Receptor* | ENSG00000135503 |
| ACVR1C | *RSTK Receptor* | ENSG00000123612 |
| ACVR2A | *RSTK Receptor* | ENSG00000121989 |
| ACVR2B | *RSTK Receptor* | ENSG00000114739 |
| ACVRL1 | *RSTK Receptor* | ENSG00000139567 |
| ADAMDEC1 | *ADAMDEC1 Mesenchyme* | ENSG00000134028 |
| ALDH1A3 | *ADAMDEC1 Mesenchyme* | ENSG00000184254 |
| ALPI | *Enterocyte* | ENSG00000163295 |
| AMHR2 | *RSTK Receptor* | ENSG00000135409 |
| ANO1 | *ICC* | ENSG00000131620 |
| AREG | *Paneth cell; Innate lymphoid cell* | ENSG00000109321 |
| ARX | *Enteroendocrine cell* | ENSG00000004848 |
| ASCL2 | *Stem cell* | ENSG00000183734 |
| ASPN | *ECM* | ENSG00000106819 |
| ATOH1 | *Early Secretory; Notch Signaling* | ENSG00000172238 |
| BANK1 | *Immune: memory B cell* | ENSG00000153064 |
| BEST2 | *Goblet cell* | ENSG00000039987 |
| BEST4 | *Paneth cell; BEST4+ Epithelial cell* | ENSG00000142959 |
| BMP2 | *BMP Signaling* | ENSG00000125845 |
| BMP3 | *BMP3+ SEC* | ENSG00000152785 |
| BMP4 | *BMP Signaling* | ENSG00000125378 |
| BMP5 | *RSTK Ligand* | ENSG00000112175 |
| BMP7 | *RSTK Ligand* | ENSG00000101144 |
| BMP8B | *RSTK Ligand* | ENSG00000116985 |
| BMPR1A | *RSTK Receptor* | ENSG00000107779 |
| BMPR2 | *RSTK Receptor* | ENSG00000204217 |
| BTC | *EGF Signaling* | ENSG00000174808 |
| C1QTNF3 | *SHISA3+ Mesenchyme* | ENSG00000082196 |
| CA4 | *Paneth cell; BEST4+ Epithelial cell* | ENSG00000167434 |
| CA7 | *Paneth cell; BEST4+ Epithelial cell* | ENSG00000168748 |
| CAPN3 | *Mesenchyme* | ENSG00000092529 |
| CCL19 | *CCL19+ Mesenchyme* | ENSG00000172724 |
| CCL2 | *CCL2/FABP5 Mesenchyme* | ENSG00000108691 |
| CCL21 | *Endothelium* | ENSG00000137077 |
| CCL5 | *Immune: gamma-delta T cell* | ENSG00000271503 |
| CD14 | *Immune: macrophage* | ENSG00000170458 |
| CD163 | *Immune: macrophage* | ENSG00000177575 |
| CD19 | *Immune: B cells (all)* | ENSG00000177455 |
| CD1D | *Immune: Breg* | ENSG00000158473 |
| CD27 | *Immune: B plasmablast* | ENSG00000139193 |
| CD34 | *Endothelial cells, MSCs* | ENSG00000174059 |
| CD38 | *Immune: B plasma cell* | ENSG00000004468 |
| CD3E | *Immune: Tregs, T helper cells, T follicular helper cells, Tfr, Th17, Th9, Th22* | ENSG00000198851 |
| CD4 | *Immune: CD4+ T cells* | ENSG00000010610 |
| CD5 | *Immune: T-helper 1 cell, B cell marker* | ENSG00000110448 |
| CD68 (SRD1) | *Immune: macrophage, mouse ILC3* | ENSG00000129226 |
| CD69 | *Immune: eosinophil, Trm* | ENSG00000110848 |
| CD7 | *Immune: gamma-delta T cell* | ENSG00000173762 |
| CD79A (IgA) | *Immune: B cell* | ENSG00000105369 |
| CD81 | *Mesenchyme* | ENSG00000110651 |
| CD8A | *Immune: CD8-positive, alpha-beta T cell* | ENSG00000153563 |
| CDH1 | *Epithelium* | ENSG00000039068 |
| CDH5 (CD144, VE-Cadherin) | *Endothelium* | ENSG00000179776 |
| CDK15 | *Immune: mast cell* | ENSG00000138395 |
| CDX2 | *Epithelium* | ENSG00000165556 |
| CEACAM6 | *Paneth cell; BEST4+ Epithelial cell* | ENSG00000086548 |
| CEACAM7 | *Enterocyte; BEST4+ Epithelial cell* | ENSG00000007306 |
| CFTR | *Progenitor cell; Absorptive cell* | ENSG00000001626 |
| CHAT | *ENS (cholinergic neurons)* | ENSG00000070748 |
| CHGA | *Enteroendocrine cell* | ENSG00000100604 |
| CHGB | *Enteroendocrine cell* | ENSG00000089199 |
| CMA1 | *Immune: mast cell* | ENSG00000092009 |
| COL14A1 | *SHISA3+ Mesenchyme* | ENSG00000187955 |
| COL16A1 | *ECM* | ENSG00000084636 |
| COL17A1 | *Enterocyte; Absorptive cell* | ENSG00000065618 |
| COL18A1 | *ECM* | ENSG00000182871 |
| COL1A1 | *Mesenchyme* | ENSG00000108821 |
| COL1A2 | *Mesenchyme* | ENSG00000164692 |
| COL21A1 | *ECM* | ENSG00000124749 |
| COL23A1 | *ECM* | ENSG00000050767 |
| COL3A1 | *ECM* | ENSG00000168542 |
| COL4A5 | *ECM* | ENSG00000188153 |
| COL6A1 | *ECM* | ENSG00000142156 |
| COL6A2 | *ECM* | ENSG00000142173 |
| COL9A1 | *ECM* | ENSG00000112280 |
| COX4I2 | *Pericyte* | ENSG00000131055 |
| CPA3 | *Immune: mast cell (granulocytes)* | ENSG00000163751 |
| CPM | *BMP3+ SEC* | ENSG00000135678 |
| CST7 | *Immune: CD8-positive, alpha-beta T cell* | ENSG00000077984 |
| CTLA4 | *Immune: Type A IEL* | ENSG00000163599 |
| CTSG | *Immune: mast cell, granulocytes* | ENSG00000100448 |
| CX3CR1 | *Immune: CX3CR1+ dendritic cell, monocyte, macrophage* | ENSG00000168329 |
| CXCL13 | *CCL19+ Mesenchyme* | ENSG00000156234 |
| CXCL14 | *NPY+ SEC and ADAMDEC1 Mesenchyme* | ENSG00000145824 |
| CXCR4 | *BMP3+ SEC* | ENSG00000121966 |
| DCN | *ECM* | ENSG00000011465 |
| DEFA5 | *Paneth cells* | ENSG00000164816 |
| DEFA6 | *Paneth cells* | ENSG00000164822 |
| DERL3 | *Immune: IgA plasma cell* | ENSG00000099958 |
| DES | *VSMC* | ENSG00000175084 |
| DHH | *HH Signaling* | ENSG00000139549 |
| DKK3 | *WNT Antagonist* | ENSG00000050165 |
| DLL1 | *Goblet cells, SEC* | ENSG00000198719 |
| DONSON | *Mesenchyme* | ENSG00000159147 |
| DPT | *Mesenchyme* | ENSG00000143196 |
| EDNRB | *Mesenchyme* | ENSG00000136160 |
| EGF | *EGF Signaling* | ENSG00000138798 |
| EGFR | *EGF Signaling* | ENSG00000146648 |
| ELAVL3 (HUC) | *ENS (generic neuron marker)* | ENSG00000196361 |
| EPCAM | *Epithelium* | ENSG00000119888 |
| EPGN | *EGF Signaling* | ENSG00000182585 |
| ERBB2 | *EGF Signaling* | ENSG00000141736 |
| ERBB3 | *EGF Signaling* | ENSG00000065361 |
| ERBB4 | *EGF Signaling* | ENSG00000178568 |
| EREG | *EGF Signaling* | ENSG00000124882 |
| ETS1 | *Immune: T-helper 1 cell* | ENSG00000134954 |
| F3 | *SEC* | ENSG00000117525 |
| FABP2 | *Enterocyte* | ENSG00000145384 |
| FABP5 | *CCL2/FABP5 Mesenchyme* | ENSG00000164687 |
| FABP7 | *Pericyte* | ENSG00000164434 |
| FBLN1 | *ECM* | ENSG00000077942 |
| FBXL22 | *Mesenchyme (between ICM and OLM)* | ENSG00000197361 |
| FCGR3A (CD16) | *Immune: neutrophil, NKreg, NKcyto* | ENSG00000203747 |
| FCRLA | *Immune: memory B cell* | ENSG00000132185 |
| FIBIN | *SHISA3+ Mesenchyme* | ENSG00000176971 |
| FN1 | *ECM* | ENSG00000115414 |
| FNDC1 | *ECM* | ENSG00000164694 |
| FOXF1 | *HH Target* | ENSG00000103241 |
| FOXL1 | *SEC* | ENSG00000176678 |
| FOXP3 | *Immune: follicular T reg cells, Treg* | ENSG00000049768 |
| FRZB | *Goblet cell, NPY+ SEC* | ENSG00000162998 |
| FST | *SMC and precursor* | ENSG00000134363 |
| FZD1 | *WNT Receptor* | ENSG00000157240 |
| FZD10 | *WNT Receptor* | ENSG00000111432 |
| FZD2 | *WNT Receptor* | ENSG00000180340 |
| FZD3 | *WNT Receptor* | ENSG00000104290 |
| FZD4 | *WNT Receptor* | ENSG00000174804 |
| FZD5 | *WNT Receptor* | ENSG00000163251 |
| FZD6 | *WNT Receptor* | ENSG00000164930 |
| FZD7 | *WNT Receptor* | ENSG00000155760 |
| FZD8 | *WNT Receptor* | ENSG00000177283 |
| GATA2 | *Immune: mast cell (pan hematopoietic marker)* | ENSG00000179348 |
| GATA3 | *Immune: Th9, ILC2* | ENSG00000107485 |
| GDF10 | *RSTK Ligand* | ENSG00000266524 |
| GDF11 | *RSTK Ligand* | ENSG00000135414 |
| GDF6 | *RSTK Ligand* | ENSG00000156466 |
| GLI1 | *HH Signaling* | ENSG00000111087 |
| GLI2 | *HH Target* | ENSG00000074047 |
| GPC3 | *ECM* | ENSG00000147257 |
| GPX3 | *ADAMDEC1 Mesenchyme* | ENSG00000211445 |
| GREM1 | *Smooth Muscle; MSCs* | ENSG00000166923 |
| GUCA2A | *Paneth, BEST4+ Epithelial cell* | ENSG00000197273 |
| GUCA2B | *Paneth, BEST4+ Epithelial cell* | ENSG00000044012 |
| GZMA | *Immune: gamma-delta T cell, NK cells* | ENSG00000145649 |
| GZMB | *Immune: Trm* | ENSG00000100453 |
| GZMK | *Immune: CD8-positive, alpha-beta T cell* | ENSG00000113088 |
| HAPLN1 | *SEC* | ENSG00000145681 |
| HBEGF | *EGF Signaling* | ENSG00000113070 |
| HDC | *Immune: Mast cell (myeloid cell and myeloid biased hematopoietic stem cells)* | ENSG00000140287 |
| HGF | *HGF Signaling* | ENSG00000019991 |
| HHIP | *NPY+ SEC, HH Signaling* | ENSG00000164161 |
| HLA-DRB1 (MHC-II) | *Immune: APCs=dendritic cells, macrophages, B cells* | ENSG00000196126 |
| HMGB2 | *Transit amplifying cell; Secretory progenitor cell* | ENSG00000164104 |
| HPGDS | *Immune: mast cell; tuft cell* | ENSG00000163106 |
| HTR3E | *Tuft cell* | ENSG00000186038 |
| ICAM1 | *Immune: FDC* | ENSG00000090339 |
| ICOS | *Immune: follicular T cells (helper and reg)* | ENSG00000163600 |
| ID1 | *NPY+ SEC, Epithelium, and Endothelium* | ENSG00000125968 |
| ID2 | *BMP Target* | ENSG00000115738 |
| IGFBP3 | *NPY+ SEC, Endothelium* | ENSG00000146674 |
| IHH | *HH Signaling* | ENSG00000163501 |
| IL10 | *Immune: macrophage, Breg, mouse ILC3* | ENSG00000136634 |
| IL12RB1 | *Immune: Th17, Trm* | ENSG00000096996 |
| IL17A | *Immune: Th17* | ENSG00000112115 |
| IL23A | *Immune: macrophage* | ENSG00000110944 |
| IL23R | *Immune: Th17, ILC3* | ENSG00000162594 |
| IL2RA (CD25) | *Immune: follicular T reg cells, T regs, ILC2* | ENSG00000134460 |
| IL6R | *Immune: Th17* | ENSG00000160712 |
| IL7R (CD127) | *Immune: T-helper 17 cell* | ENSG00000168685 |
| INHBA | *RSTK Ligand* | ENSG00000122641 |
| ITGAL (CD11a) | *Immune: type A IEL* | ENSG00000005844 |
| ITGAM (CD11b) | *Immune: eosinophil, monocyte, neutrophil* | ENSG00000169896 |
| ITGB2 (CD18) | *Immune: eosinophil, neutrophil* | ENSG00000160255 |
| KIT | *Immune: mast cell, ICC* | ENSG00000157404 |
| KLRB1 | *Immune: T-helper 1 cell, Th17 cells* | ENSG00000111796 |
| KREMEN1 | *WNT Interacting* | ENSG00000183762 |
| KRT19 | *Mesothelium, Epithelium* | ENSG00000171345 |
| LGALS2 | *Progenitor cell; dendritic cell; absorptive cell* | ENSG00000100079 |
| LGR4 | *WNT Interacting* | ENSG00000205213 |
| LGR5 | *Stem cell* | ENSG00000139292 |
| LRP5 | *WNT Interacting* | ENSG00000162337 |
| LRP6 | *WNT Interacting* | ENSG00000070018 |
| LTB | *Immune: regulatory T cell* | ENSG00000227507 |
| LUM | *ECM* | ENSG00000139329 |
| LYVE1 | *Immune: macrophage, Lymphatic endothelial cell* | ENSG00000133800 |
| LYZ | *Paneth, M cell, FAE, BEST4+* | ENSG00000090382 |
| MEIS2 | *Immune: mast cell, mesenchyme, and endothelium* | ENSG00000134138 |
| MET | *HGF Signaling* | ENSG00000105976 |
| MKI67 | *Transit amplifying cell; dendritic cell* | ENSG00000148773 |
| MMP11 | *BMP3+ SEC* | ENSG00000099953 |
| MPPED2 | *OLM and serosa* | ENSG00000066382 |
| MS4A1 (CD20) | *Immune: memory B cell* | ENSG00000156738 |
| MS4A2 | *Immune: mast cell (granulocytes)* | ENSG00000149534 |
| MSLN | *Mesothelium* | ENSG00000102854 |
| MSTN | *RSTK Ligand* | ENSG00000138379 |
| MUC2 | *Goblet cells* | ENSG00000198788 |
| MYH11 | *Myofibroblasts / Myo-like* | ENSG00000133392 |
| MYLK | *Myofibroblasts / Myo-like* | ENSG00000065534 |
| NDP | *WNT Interacting* | ENSG00000124479 |
| NEUROD1 | *Enteroendocrine cell* | ENSG00000162992 |
| NKX2-3 | *Mesenchyme* | ENSG00000119919 |
| NPY | *NPY+, SEC* | ENSG00000122585 |
| NRG1 | *EGF Signaling* | ENSG00000157168 |
| NRG2 | *EGF Signaling* | ENSG00000158458 |
| NRG3 | *EGF Signaling* | ENSG00000185737 |
| NRG4 | *EGF Signaling* | ENSG00000169752 |
| NRP2 | *Mesenchyme* | ENSG00000118257 |
| ODF2L | *Immune: T-helper 17 cell* | ENSG00000122417 |
| OGN | *ECM* | ENSG00000106809 |
| OLFM4 | *Transit amplifying cell* | ENSG00000102837 |
| OSR1 | *Mesenchyme* | ENSG00000143867 |
| OTOP2 | *Paneth cell; BEST4+ epithelial cell* | ENSG00000183034 |
| PAX4 | *Enteroendocrine cell* | ENSG00000106331 |
| PAX5 | *Immune: memory B cell* | ENSG00000196092 |
| PCNA | *Proliferation* | ENSG00000132646 |
| PDGFRA | *Mesenchyme, SEC* | ENSG00000134853 |
| PDGFRB | *Mesenchyme* | ENSG00000113721 |
| PDX1 | *Proximal small intestine epithelium* | ENSG00000139515 |
| PECAM1 (CD31) | *Endothelium* | ENSG00000261371 |
| PROX1 | *Lymphatic endothelial cell* | ENSG00000117707 |
| PTCH1 | *HH Signaling* | ENSG00000185920 |
| PTGS1 (COX1) | *Human tuft cell marker* | ENSG00000095303 |
| PTPRC (CD45) | *Immune marker* | ENSG00000081237 |
| RBP2 | *Enterocyte* | ENSG00000114113 |
| RBP5 | *CCL19+ mesenchyme, endothelium* | ENSG00000139194 |
| REG4 | *Goblet cell* | ENSG00000134193 |
| RFX6 | *Enteroendocrine cell* | ENSG00000185002 |
| RGS5 | *VSMC* | ENSG00000143248 |
| RNF43 | *Stem cell* | ENSG00000108375 |
| RRM2 | *Transit amplifying cell; Secretory progenitor cell* | ENSG00000171848 |
| RSPO2 | *WNT Interacting* | ENSG00000147655 |
| RSPO3 | *WNT Interacting* | ENSG00000146374 |
| S100B | *ENS (Glia)* | ENSG00000160307 |
| SATB2 | *Distal small intestine epithelium* | ENSG00000119042 |
| SELL (CD62L) | *Immune: neutrophil, Trm (low)* | ENSG00000188404 |
| SFRP1 | *SFRP1+ proliferative mesenchyme* | ENSG00000104332 |
| SFRP2 | *ECM* | ENSG00000145423 |
| SHH | *HH Signaling* | ENSG00000164690 |
| SHISA3 | *SHISA3+ Mesenchyme* | ENSG00000178343 |
| SI | *Epithelium enterocytes* | ENSG00000090402 |
| SLC12A2 | *Stem cell* | ENSG00000064651 |
| SLC18A2 | *Immune: mast cell (granulocytes)* | ENSG00000165646 |
| SMAD4 | *SMAD (co-mediated)* | ENSG00000141646 |
| SMO | *WNT Interacting* | ENSG00000128602 |
| SPARCL1 | *ECM* | ENSG00000152583 |
| SPDEF | *Goblet cells* | ENSG00000124664 |
| SPIB | *Paneth cell; BEST4+ Epithelial cell* | ENSG00000269404 |
| SPOCK2 | *Immune: T-helper 17 cell* | ENSG00000107742 |
| STMN1 | *Transit amplifying cell; Dendritic cell; Secretory progenitor cell* | ENSG00000117632 |
| TAGLN | *Myofibroblasts / Myo-like* | ENSG00000149591 |
| TAS1R3 | *Tuft cell* | ENSG00000169962 |
| TBX21 (T-bet) | *Immune: Th22, mouse ILC3* | ENSG00000073861 |
| TFF1 | *Goblet cell* | ENSG00000160182 |
| TFF3 | *Goblet cells* | ENSG00000160180 |
| TGFA | *EGF Signaling* | ENSG00000163235 |
| TGFB1 | *RSTK Ligand* | ENSG00000105329 |
| TGFB2 | *RSTK Ligand* | ENSG00000092969 |
| TGFB3 | *RSTK Ligand* | ENSG00000119699 |
| TGFBR1 | *RSTK Receptor* | ENSG00000106799 |
| TGFBR2 | *RSTK Receptor* | ENSG00000163513 |
| TGFBR3 | *RSTK Receptor* | ENSG00000069702 |
| THY1 (CD90) | *Mesenchyme* | ENSG00000154096 |
| TIGIT | *Immune: regulatory T cell* | ENSG00000181847 |
| TK1 | *Transit amplifying cell; Secretory progenitor cell* | ENSG00000167900 |
| TNFAIP3 | *Immune: T-helper 17 cell* | ENSG00000118503 |
| TNFRSF17 | *Immune: IgA plasma cell* | ENSG00000048462 |
| TNFRSF18 | *Immune: Follicular T regs, T regs, B plasmablast* | ENSG00000186891 |
| TOP2A | *Proliferation* | ENSG00000131747 |
| TPH1 | *ENS (serotonergic neurons)* | ENSG00000129167 |
| TRPM5 | *Tuft cell* | ENSG00000070985 |
| TUBB3 | *ENS (neuron + glia marker)* | ENSG00000258947 |
| TYMS | *Transit amplifying cell; Dendritic cell* | ENSG00000176890 |
| UBE2C | *Transit amplifying cell; Dendritic cell; Secretory progenitor cell* | ENSG00000175063 |
| UCHL1 (PGP9.5) | *ENS, generic neuron marker* | ENSG00000154277 |
| UPK3B | *Mesothelium* | ENSG00000243566 |
| VCAM1 | *Immune: FDC* | ENSG00000162692 |
| VIL1 | *Epithelium* | ENSG00000127831 |
| VIM | *Mesenchyme* | ENSG00000026025 |
| VIP | *ENS (VIPergic neurons)* | ENSG00000146469 |
| VPREB3 | *Immune: memory B cell* | ENSG00000128218 |
| WIF1 | *WNT Interacting* | ENSG00000156076 |
| WNT11 | *WNT Ligand* | ENSG00000085741 |
| WNT2B | *WNT Ligand* | ENSG00000134245 |
| WNT3 | *WNT Ligand* | ENSG00000108379 |
| WNT4 | *WNT Ligand* | ENSG00000162552 |
| WNT5A | *WNT Ligand* | ENSG00000114251 |
| WNT5B | *WNT Ligand* | ENSG00000111186 |
| WNT9A | *WNT Ligand* | ENSG00000143816 |
| WT1 | *Mesothelium* | ENSG00000184937 |

|  | **scRNAseq:**  **Major Cell Classes** | **Xenium:**  **Major Cell Classes** |
| --- | --- | --- |
| **Epithelium** | *CDH1, EPCAM, CDX2* | *CDH1, EPCAM, CDX2* |
| **Fibroblast** | *VIM, COL1A1, COL1A2, THY1, CDH2* | *VIM, THY1* |
| **SMC** | *ACTA2, TAGLN* | *ACTA2, TAGLN* |
| **ENS** | *TUBB3, ELAVL4, UCHL1, S100β, CHAT, VIP* | *TUBB3, UCHL1, S100β, CHAT, VIP, TPH1* |
| **Endothelium** | *PECAM1, CDH5* | *PECAM1, CDH5* |
| **Immune** | *PTPRC (CD45)* | *PTPRC (CD45)* |

**Supplemental Table 3:** Markers used to annotate the major cell classes in the scRNAseq and Xenium datasets

|  | **scRNAseq:**  **Fibroblast Classes** | **Xenium:**  **Fibroblast Classes** |
| --- | --- | --- |
| **Fibroblast** | *VIM, COL1A1, COL1A2, THY1, CDH2* | *VIM, THY1* |
| **Proliferative** | *MKI67, PCNA, TOP2A* | *MKI67, PCNA, TOP2A, SFRP1* |
| **SMC** | *ACTA2, TAGLN* | *ACTA2, TAGLN* |
| **VSMC/pericytes** | *DES, RGS5*, *KCNL8, ABCC9, BGN^hi^, CCDC3^hi^* | *RGS5, DES, FABP5* |
| **ICC** | *KIT, ANO1* | *KIT* |
| **SEC** | *F3, PDGFRA* | *F3, PDGFRA* |
| **LPF** | *FAPB5, ADAMDEC1, ALDH1A3, GPX3* | *FABP5, ADAMDEC1* |
| **SMF** | *SHISA3, FIBIN, C1QTNF3, COL14A1, FBLN1* | *SHISA3, FIBIN, C1QTNF3* |
| ***CXCL13*+ Fibroblast** | *CXCL13, CCL19* | *CXCL13* |

**Supplemental Table 4:** Markers used to annotate the fibroblast classes in the scRNAseq and Xenium datasets
