## Supplemental Figures 1-4 for "Mapping mesenchymal diversity in the developing human intestine and organoids"

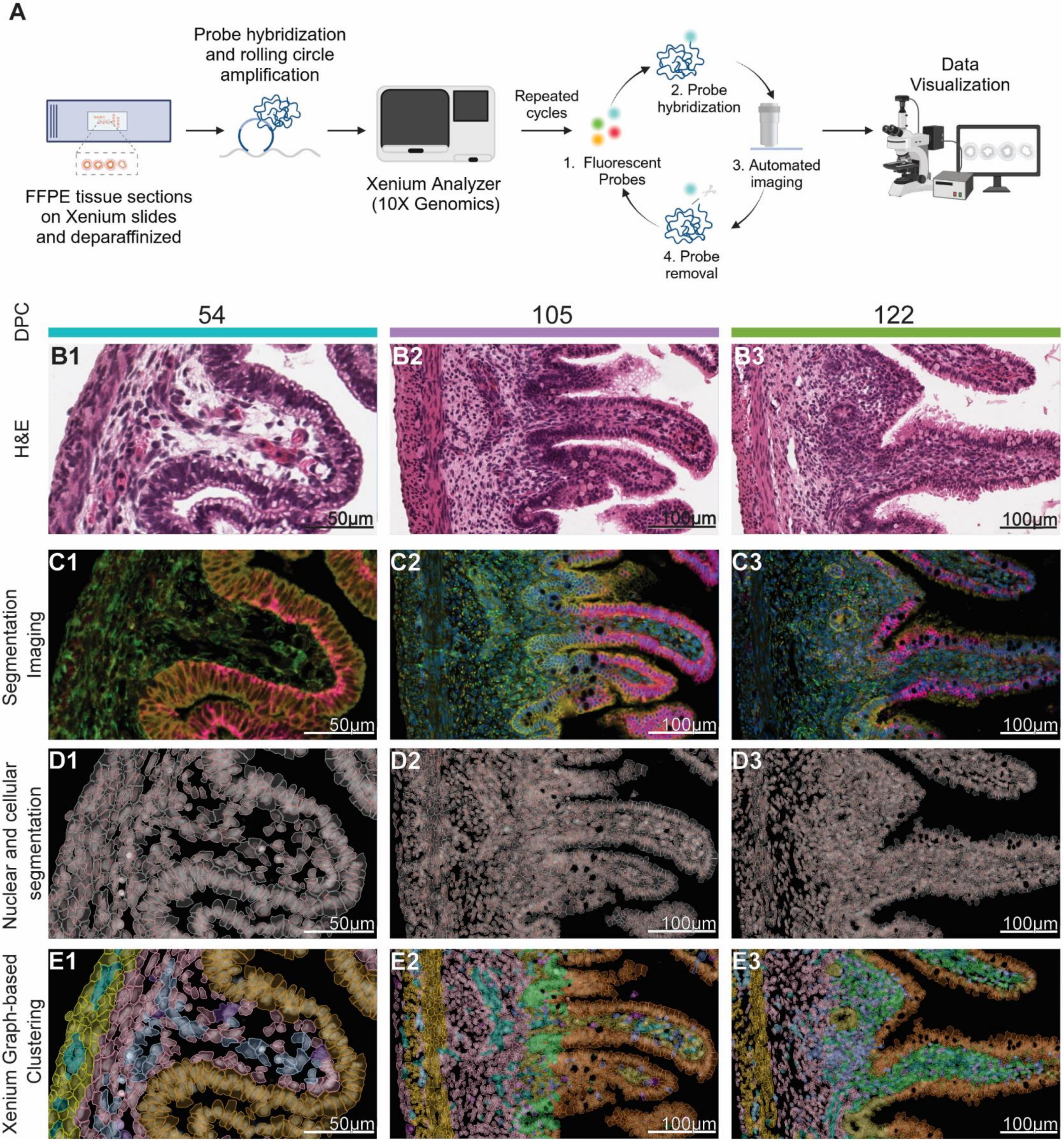


**Supplemental Figure 1: Xenium spatial transcriptomics workflow**

**A)** Workflow overview for Xenium gene expression assay. Representative Xenium images of the duodenum showing **B)** H&E, **C)** segmentation staining (PN-1000661), **D)** nuclear and cellular segmentation determined digitally using Xenium Explorer v2.0.0+ and the segmentation kit showed in **C**, and **E)** example Xenium cell clustering with assigned K equal to UMAP identified and confirmed populations.

Scale bars for images in **column 1**) 50µm and **columns 2-3)** 100µm. Colored bars indicate the developmental age group of each sample (DPC).

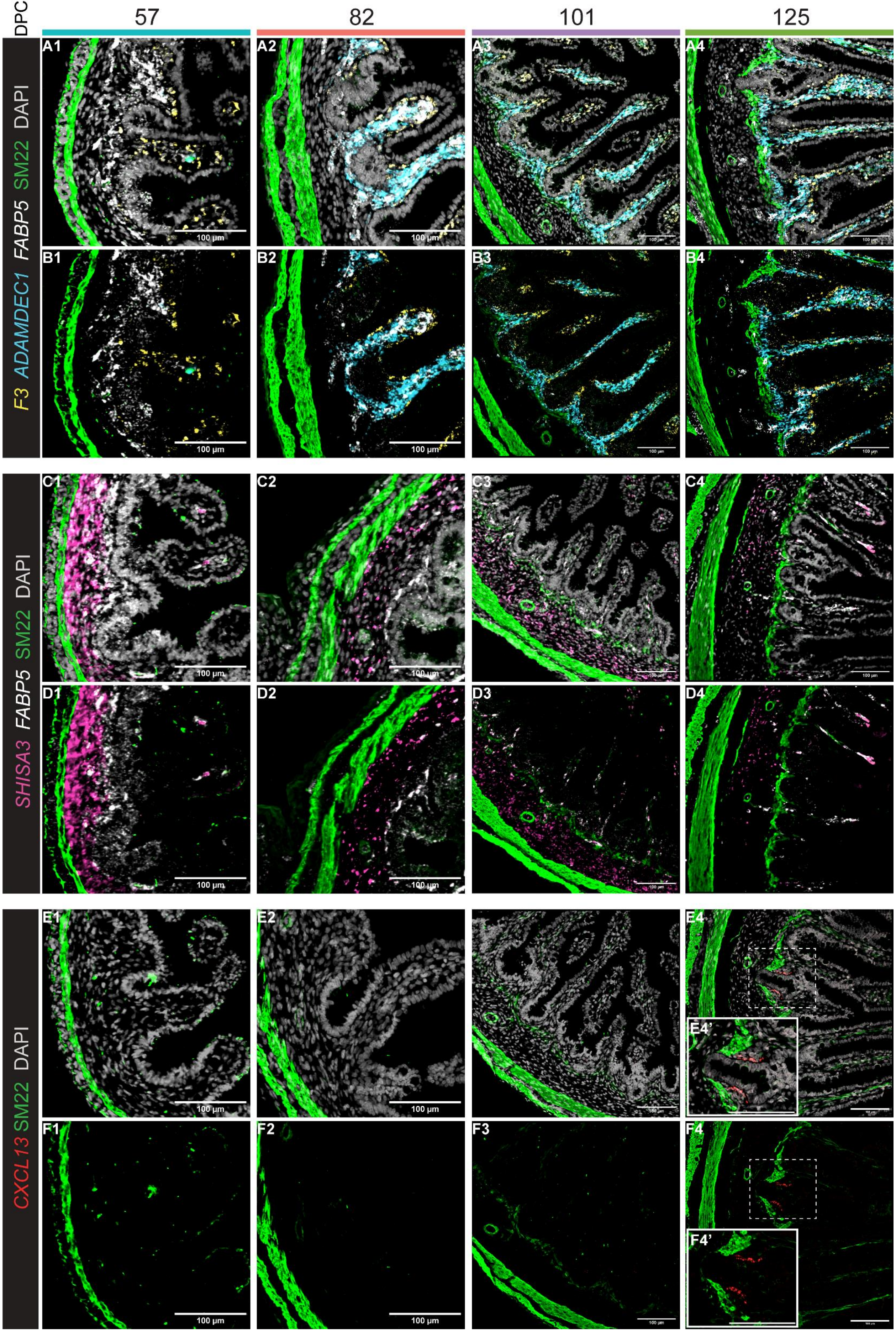


**Supplemental Figure 2: FISH validation of mesenchymal marker gene localization**

FISH staining for proposed markers of the fibroblast populations in the developing duodenum: **A-B)** SEC marker *F3* (yellow), *ADAMDEC1* (cyan), *FABP5* (white). **C-D)** *SHISA3* (pink), *FABP5* (white), **E-F)** *CXCL13* (red). DAPI (white) in images **A, C,** and **E**.
SMCs visualized with either αSMA (57 DPC) or SM22 (82-125 DPC) (green) in all images.
Scale bars = 100um.
Colored bars indicate the developmental stage (DPC) each sample falls within.

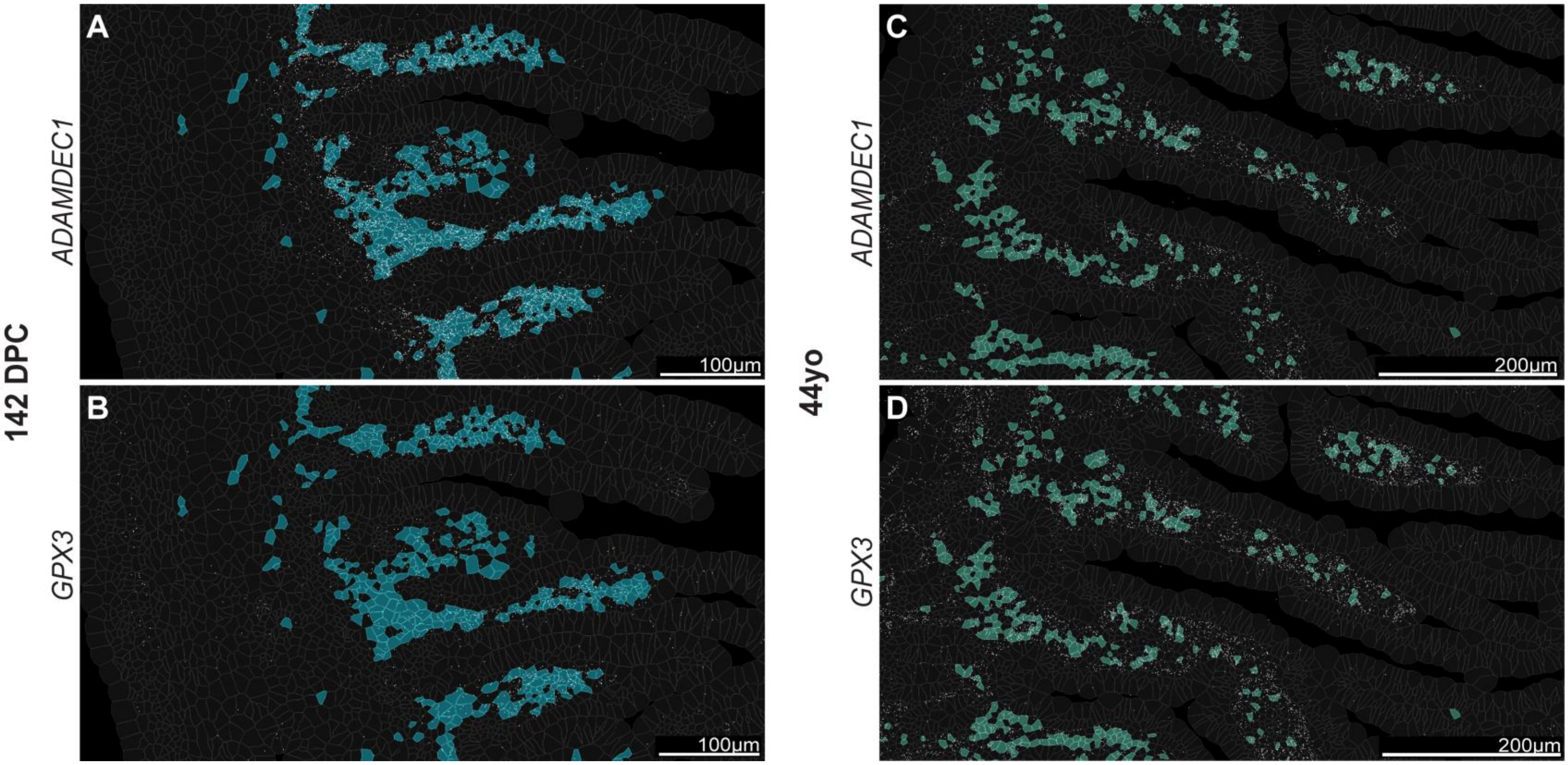


**Supplemental Figure 3: Temporal expression of LPF marker genes *ADAMDEC1* and *GPX3***

LPF cell masks for **A-B)** fetal and **C-D)** adult tissue highlighting the difference in expression of the genes *ADAMDEC1* and *GPX3.* **A)** *ADAMDEC1* is evenly expressed across all LPFs in fetal tissue (142 DPC) while **B)** *GPX3* expression is relatively lower in the fetal duodenum with a gradient along the villus, with *GPX3* being expressed highly at the villus tip while absent from the villus shelf/crypt. In the adult **C)** *ADAMDEC1* expression is relatively lower than **D)** *GPX3* expression and *GPX3* has lost the gradient of expression and is expressed throughout the entire lamina propria compartment.

Scale bars for images **A-B)** 100µm and **C-D)** 200µm.

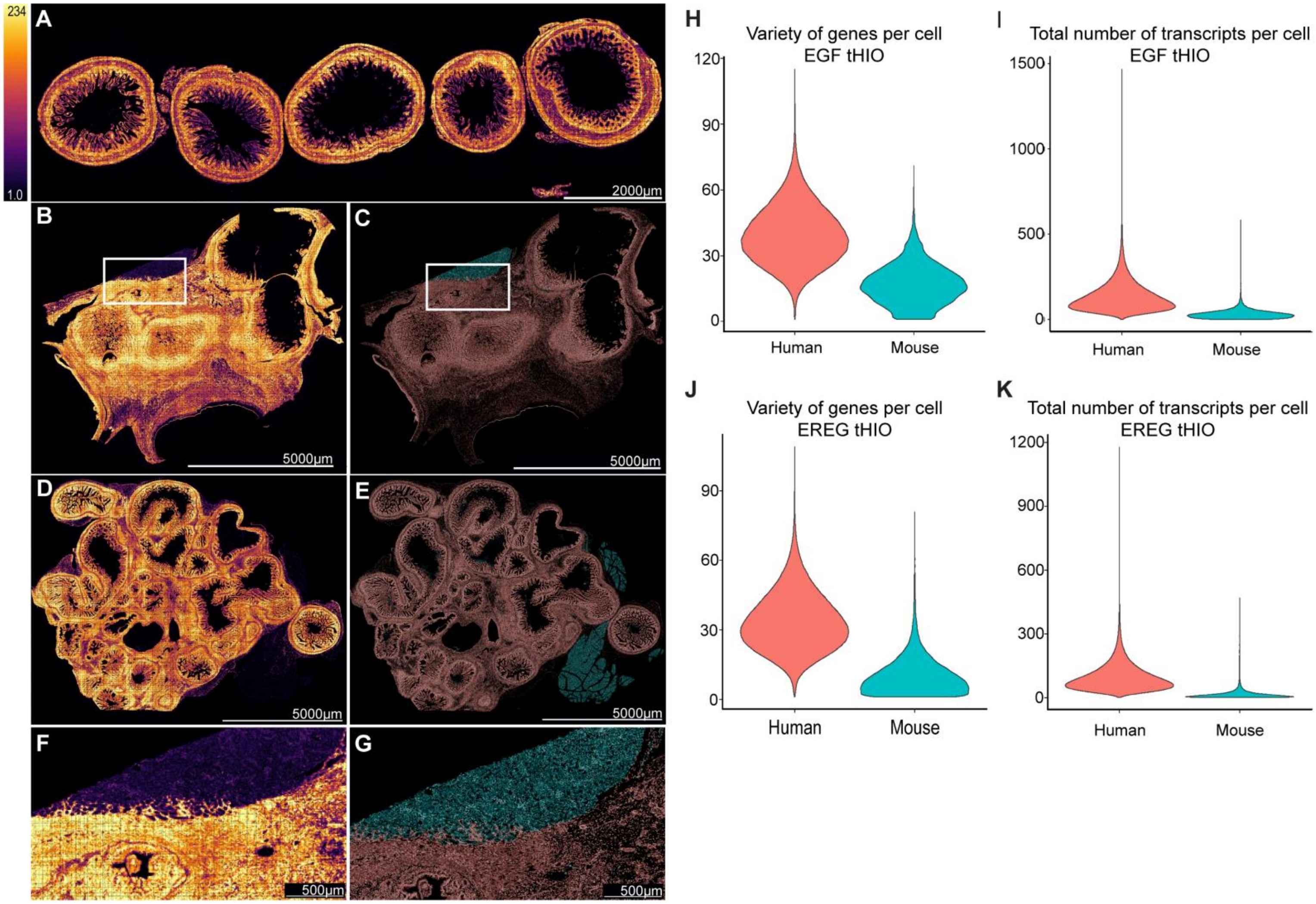


**Supplemental Figure 4: Identification of mouse cells in tHIOs**

Xenium gene panels exhibit species-specific binding. A heatmap of total transcript density across all probes for **A)** human fetal tissue (142 DPC), **B)** EGF-derived tHIO, and **D)** EREG-derived tHIO. Xenium clustering analysis identifies and clusters low-transcript expressing cells separately from high transcript expressing cells in **C)** EGF-derived tHIOs and **E)** EREG-derived tHIOs. High-magnification **F)** heatmap and **G)** clustering of the ROI depicted in **B** and **C** to highlight infiltrating human cells in the mouse tissue that would be excluded from tHIO analysis if cells were excluded on the basis of ROI selection. Violin plots of the distribution of **H, J)** the variety of transcripts and **I, H)** the total number of transcripts identified in cells of the mouse (low-transcript clusters) and human (high-transcript clusters) in the **H, I)** EGF-derived and **J, K)** EREG-derived tHIOs. Scalebars for full scale images = 5000µm, for F-G scalebars = 500µm. Human-annotated cells are highlighted and displayed in red, mouse annotated cells highlighted and displayed in blue
